# A mechanistic theory of planning in prefrontal cortex

**DOI:** 10.1101/2025.09.23.677709

**Authors:** Kristopher T. Jensen, Peter Doohan, Mathias Sablé-Meyer, Sandra Reinert, Alon Baram, Maneesh Sahani, Thomas Akam, Timothy E. J. Behrens

## Abstract

Planning is critical for adaptive behaviour in a changing world, because it lets us anticipate the future and adjust our actions accordingly. While prefrontal cortex is crucial for this process, it remains unknown how planning is implemented in neural circuits. Prefrontal representations were recently discovered in simpler sequence memory tasks, where different populations of neurons represent different future time points. We demonstrate that combining such representations with the ubiquitous principle of neural attractor dynamics allows circuits to solve much richer problems including planning. This is achieved by embedding the environment structure directly in synaptic connections to implement an attractor network that infers desirable futures. The resulting ‘spacetime attractor’ excels at planning in challenging tasks known to depend on prefrontal cortex. Recurrent neural networks trained by gradient descent on such tasks learn a solution that precisely recapitulates the spacetime attractor – in representation, in dynamics, and in connectivity. Analyses of networks trained across different environment structures reveal a generalisation mechanism that rapidly reconfigures the world model used for planning, without the need for synaptic plasticity. The spacetime attractor is a testable mechanistic theory of planning. If true, it would provide a path towards detailed mechanistic understanding of how prefrontal cortex structures adaptive behaviour.

## Introduction

While different cortical areas support different functions, common computational principles are shared across many areas. For functions as disparate as sensory processing, spatial reasoning, and language comprehension, features of the environment are inferred from partial information. To do so, structural knowledge about the world must be embedded in synaptic connections (Ko et al., 2011; Burak and Fiete, 2009; Iacaruso et al., 2017; Turner-Evans et al., 2020). This constrains neural circuits to represent meaningful interpretations of the environment, and inputs select between these interpretations. It is not known whether similar principles generalise to complex prospective behaviours, such as planning extended action sequences to achieve a distant goal. Recent recordings from prefrontal cortex give an important clue. When mice or monkeys must execute a sequence of actions, neurons represent the entire sequence concurrently (El-Gaby et al., 2023; Xie et al., 2022). Different neuronal populations encode different steps of the future behaviour. If such a representation could be inferred from inputs indicating goals and constraints, the network could plan the future. Excitingly, this solution would use algorithmic principles similar to those known to infer features of the present in other cortical areas.

This paper has four principal aims. (i) To develop a detailed circuit model that infers explicit representations of the future from partial inputs. (ii) To discern the principles of synaptic connectivity that enable such a model to solve complex problems including planning. (iii) To understand when and why this algorithm succeeds while simpler circuit models fail. (iv) To explore how it relates to pre-frontal representations, connectivity, and function. To this end, we first construct a neural network that combines known PFC representations with structured connectivity to plan the future. We then show that the same connectivity emerges when PFC-like representations are imposed, but the weights are optimised for planning. Finally, this solution is learned by unconstrained neural networks, when both their representations and weights are optimised for challenging planning problems.

### Sequence representations in PFC

Recent work has uncovered PFC representations underlying sequence working memory (Xie et al., 2022; El-Gaby et al., 2023; Whittington et al., 2023; Botvinick and Plaut, 2006). Separate neural ‘subspaces’, or groups of neurons, represent the expected or desired state of the world at different steps along the sequence of future behaviour. In other words, some neural subspace ‘A’ represents the present, while subspace ‘B’ represents the immediate future, and subspace ‘C’ the more distant future (Figure 1A). Critically, these subspaces are active simultaneously. Together, they instantaneously represent the entire behavioural sequence. Importantly, El-Gaby et al. (2023) showed that the PFC subspaces representing different steps of the future are not independent. Neuronal correlations reflect the structure of the task being performed, even during rest.

**Figure 1.**
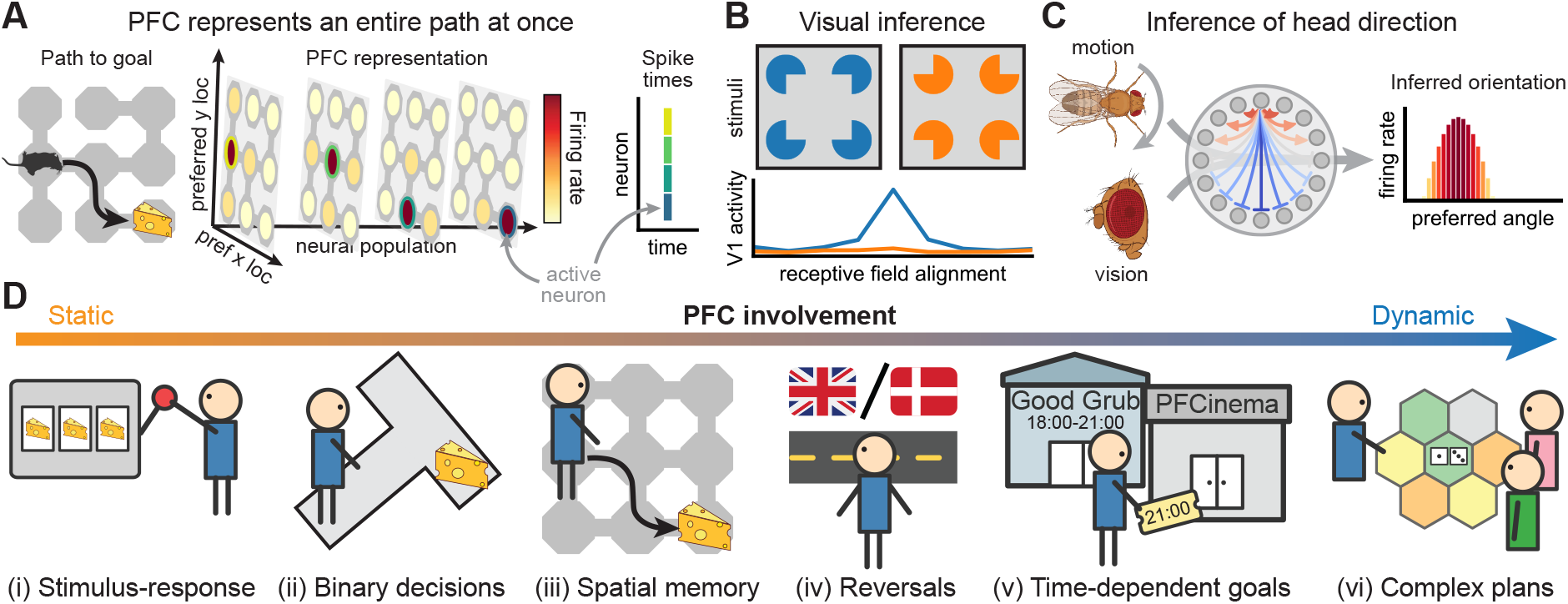
Background. **(A)** Recent work has identified explicit sequence representations in prefrontal cortex during working memory (Xie et al., 2022; El-Gaby et al., 2023). When an animal has to execute a behavioural sequence (left), individual neurons represent conjunctions of location and sequence element (centre). Separate populations (planes) therefore represent the expected location at different times in the future. The entire sequence is represented concurrently by the simultaneous firing of different neurons that encode the expected location at each time in the future (right). **(B)** An example V1 cell fires when its receptive field is aligned with an inferred line (blue), but not for a control stimulus with no inference (orange). Such visual inference is mediated by structural priors embedded in the circuit connectivity, where neurons representing consistent visual features excite each other (Iacaruso et al., 2017; Shin et al., 2023). Figure adapted from Lee and Nguyen (2001). **(C)** Structural knowledge is embedded in the synapses of the head direction system (centre; Turner-Evans et al., 2020), which constrains the network to represent single angles (Kim et al., 2017). Visual and proprioceptive inputs (left) determine which angle should be represented (right). We suggest that a mechanism like (B) and (C) infers the representation in (A). **(D)** Prefrontal cortex is particularly important in dynamic environments. Stimulus-response associations and repeated choices are robust to prefrontal lesions (i; ii), and acortical mice can solve spatial memory tasks (iii; Zheng et al., 2024). However, PFC is needed for reversal learning (iv; Walton et al., 2010) and when different goals are important, or ‘rewarding’, at different times (v; Shallice and Burgess, 1991). In multiplayer board games (vi), different resources are valuable at different stages, and opponents can dynamically change their strategy.

### Planning by inferring the future

The correlation structure observed in PFC resembles biological attractor circuits that infer the current state of the world, such as the visual input or head direction of an animal (Figure 1B-C; Ko et al., 2011; Chaudhuri et al., 2019). If similar dynamics acted on the future representation in PFC, it would extend its function beyond sequence memory to situations where entire action sequences are inferred. Planning could be solved by inferring sequences of desired actions from goals and constraints in a recurrent network implementation of ‘planning-as-inference’ (Botvinick and Toussaint, 2012; Levine, 2018; George et al., 2021). The resulting algorithm would naturally cope with dynamic environments, because it represents each time in the future separately. This is intriguing because PFC is particularly important for problems that require the correct behaviour to be expressed at an appropriate time (Figure 1D; Shallice and Burgess, 1991; Volle et al., 2011). Planning via attractor dynamics is also consistent with winnertake-all dynamics identified in frontal cortex during non-sequential behaviours (Ruff et al., 2025; Inagaki et al., 2019). Such planning-as-inference differs from most planning algorithms studied in cognitive science and machine learning, which often rely on sequential search (Callaway et al., 2022; Schrittwieser et al., 2020). Search is easily adapted to new environments, but it is slow at decision time. In familiar environments, where the structure can be embedded in cortical connections, attractor dynamics provide a complementary mechanism for rapid evaluation of many possible futures in parallel.

### A mechanistic theory

In this paper, we show that known features of PFC representations and connectivity are sufficient to implement a powerful planning algorithm with minimal additional assumptions. The resulting ‘spacetime attractor’ (STA) instantiates an explicit world model in the synaptic connections between neurons, which allows it to plan by inferring optimal future trajectories. This algorithm resembles other cortical circuits known to infer features of the present, thereby unifying our understanding of PFC with the rest of cortex. The spacetime attractor excels at dynamic problems with changing reward and transition structures, which PFC is critical for and existing mechanistic models cannot solve. RNNs trained on dynamic tasks implement a spacetime attractor in their dynamics, suggesting that it is an efficient solution. Our findings provide a precise mechanistic theory of adaptive behaviour that reconciles prior work on PFC representations with other known cortical computations and deficits from lesions.

## Results

### An attractor network in space and time

We now introduce the spacetime attractor in more detail. We first review ring and grid attractors, which illustrate how structured connectivity can guide inference from partial information. We then explain how the spacetime attractor uses these principles to infer future behaviour. Neurons are assumed to encode single environment features, but similar ideas apply when each neuron represents a combination of variables (Clark et al., 2025).

#### Ring and grid attractors

Ring attractors are neural networks that can only encode circular variables (Zhang, 1996; Ben-Yishai et al., 1995). Different neurons have different ‘preferred orientations’, and the network dynamics have a set of fixed points (stable activity patterns) that encode a particular angle. At these fixed points, neurons are only active if their preferred angle is close to the encoded angle (Figure 2A, bottom left). This is achieved by a network where neurons are mutually excitatory if they have similar preferred orientations, and inhibitory otherwise (Figure 2A, top left). The network dynamics infer which angle is most compatible with noisy inputs such as vision and proprioception (Turner-Evans et al., 2017; Skaggs et al., 1994; Kim et al., 2019; Kutschireiter et al., 2023).

**Figure 2.**
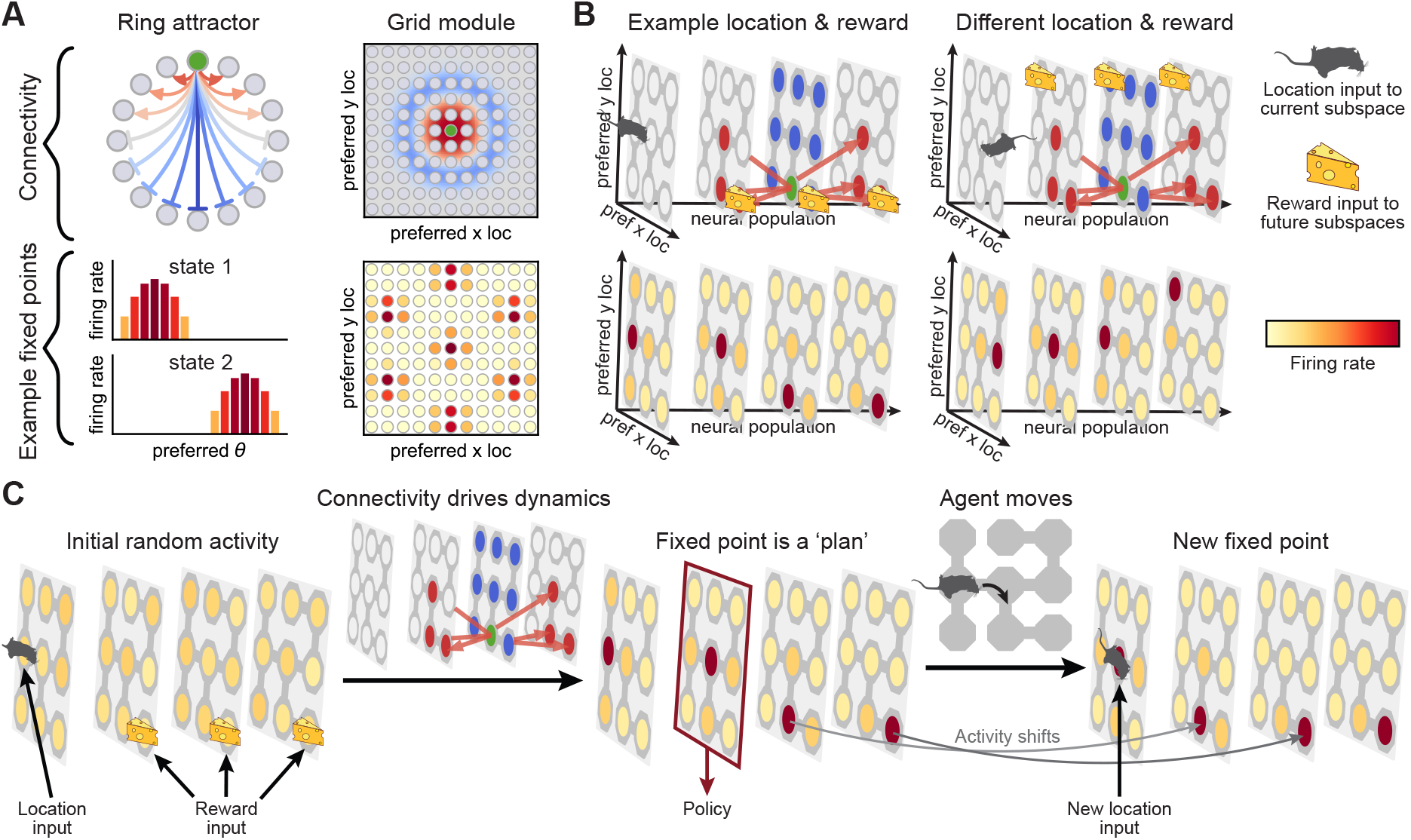
The spacetime attractor. **(A)** Left; in a one-dimensional ring attractor, neurons representing similar directions excite each other (top, red), and different directions inhibit each other (blue). Connections are shown for the green example cell. Stable network states (fixed points) have neurons at a particular encoded angle being most active (bottom). Right; grid cells (bottom) emerge from a two-dimensional attractor network (top), where cells with similar preferred locations excite each other (red) and intermediate distances inhibit each other (blue). **(B)** The spacetime attractor encodes a three-dimensional representation, where neurons have both a preferred location (planes) and delay (horizontal axis). This resembles the PFC representation in Figure 1A. Cells that represent adjacent locations in both space and time excite each other (top, red), and different locations at the same time inhibit each other (blue). Given inputs indicating the current location (mouse) and future reward (cheese), the fixed points represent reward-maximising paths through space and time (bottom). While this example has a single static goal, reward inputs can also differ between subspaces if the reward is dynamic. **(C)** Initial activity is diffuse (left), followed by convergence to a stable representation of a plan (centre). At convergence, a policy is read out from the subspace that represents the next location along the desired trajectory. When the agent moves to the next state, neural activity updates to represent the remaining plan-to-go (right; El-Gaby et al., 2023).

Grid attractors instead encode two-dimensional locations. This is achieved by neurons that are mutually excitatory if they represent nearby locations and inhibitory if they represent more distant locations (Figure 2A, top right; Fuhs and Touretzky, 2006; Burak and Fiete, 2009). The fixed points of the network are hexagonal patterns of activity that resemble canonical grid cells (Figure 2A, bottom right; Hafting et al., 2005). Inputs to the network again determine which location is represented in a particular context (Burak and Fiete, 2009).

#### Spacetime attractors

Inspired by these known biological attractor networks, we propose the existence of a *spacetime attractor* in prefrontal cortex. The STA is a network that infers explicit representations of the future, and its fixed points represent entire paths through space and time (Figure 2B, bottom; Methods). Similar to ring and grid attractors, this is achieved by connecting STA neurons such that *consistent* states excite each other and *inconsistent* states inhibit each other. In the spacetime attractor, each neuron has both a preferred location and a preferred delay *δ*, which determines how far into the future its tuning curve is defined. A neuron with *δ* = 0 fires when the agent is currently at the preferred location of that neuron. A neuron with *δ* = 3 fires when the agent expects to be at the preferred location after three actions. In such a network, consistent states can form part of a single trajectory, while inconsistent states cannot. The connectivity between neural populations with preferred delays *δ* and *δ* + 1 should therefore correspond to the structure – more specifically the adjacency matrix – of the environment (Figure 2B, top). Unlike ring and grid attractors, the fixed points of the spacetime attractor depend on tonic inputs. However, the connectivity constrains the fixed points to represent continuous trajectories for any combination of inputs. In this section, we show this empirically. Later, we will see that such connectivity is optimal for planning-as-inference.

#### Reward inputs select a trajectory

To compute a plan, it is necessary to know which states will be rewarding in the future. This reward information is provided as an input to the STA and enables fast adaptation without rewiring the synaptic connections. It alters the fixed points of the recurrent dynamics to only include trajectories that are also associated with high cumulative reward (Figure 3; Methods). This resembles a previous proposal that prefrontal cortex can compute stable ‘attraction paths’ from the current state to a distant goal (Duncan, 1990). Importantly, the STA can accommodate problems with time-varying reward structures known to engage PFC (Figure 1D; Shallice and Burgess, 1991; Volle et al., 2011; Carlesimo et al., 2014). This is possible because neural populations that represent different times in the future can receive different inputs. Given a restaurant booking in 2 hours and a cinema ticket in 4 hours, the *δ* = 2 neurons would receive reward input at the restaurant, and the *δ* = 4 neurons at the cinema. The network dynamics would then infer a future trajectory, where the agent is at each location at the appropriate time. There are several possible sources of reward input to a spacetime attractor in PFC (Supplementary Note). We focus on planning under a known reward function and therefore assume access to ground-truth rewards.

**Figure 3.**
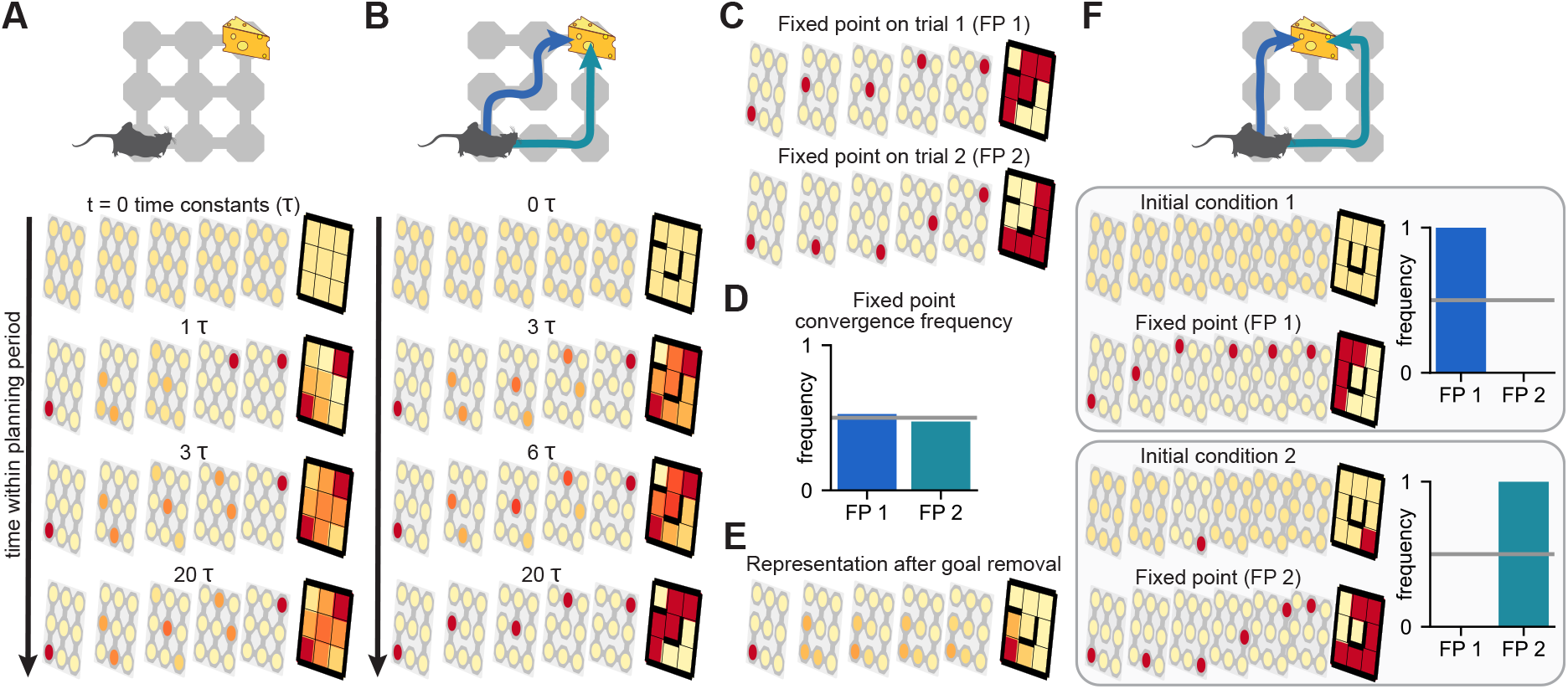
STA dynamics and fixed points. **(A)** STA dynamics on a 3 *×* 3 grid with a single goal (top). The final plane in each row is a max projection that summarises the encoded path. Time is measured in units of the network time constant *τ*. The converged representation is a stable fixed point that resembles a posterior distribution over trajectories conditioned on reaching the goal (Methods). **(B)** In an environment with several disjoint paths to the goal, there is initial competition between representations of the alternatives. The factorised representation then drives convergence to a single trajectory (Methods). **(C)** The STA converges to different fixed points on different trials in the environment from (B). **(D)** There is not a strong bias towards either of the two fixed points. **(E)** If the reward input is set to 0 after convergence in (B), the dynamics relax back to a ‘diffusive’ representation around the initial location. This illustrates the input-dependence of the fixed points. **(F)** In an environment with both a shorter and a longer path to the goal (top), the STA reliably converges to a representation of the shorter path (centre). This is because the shorter path has a higher likelihood and larger basin of attraction in the network dynamics (Methods). If the network is initialised close to a representation of the longer path, it converges to this solution instead (bottom).

To summarise, the spacetime attractor has four main components: (i) different neural populations represent different times in the future; (ii) the neurons are connected according to the structure of the environment; (iii) the current location is an input to the ‘present’ subspace; and (iv) the reward structure is an input to all future subspaces. The resulting fixed points (Figure 2B, bottom) represent trajectories that maximise cumulative reward, and they resemble an approximate posterior over the future in a probabilistic formulation of planning (Methods). The network dynamics infer future locations through a gradual relaxation process, where reward inputs bias the representation in each subspace towards high-reward locations, while inputs from neighbouring subspaces constrain their representations to be consistent (Figure 2C; Figure 3).

#### The STA guides behaviour

After computing a plan, it can be used to inform behaviour. In particular, an agent should take an action that leads to the next location on the inferred trajectory (Figure 2C, middle). The entire representation of the future then moves by one ‘action’, so the state that was previously represented by subspace *δ* is now in subspace *δ* − 1. The resulting dynamics resemble a ‘conveyor belt’ that allows the STA to execute entire trajectories without recomputing the policy. Such conveyor belt dynamics have been observed experimentally during sequence working memory (El-Gaby et al., 2023). Importantly, the representation remains a fixed point of the spacetime attractor dynamics, because the location and reward inputs are updated when the agent moves and time passes.

#### World models for planning

Many planning algorithms apply a ‘world model’ sequentially to simulate the consequences of different actions one at a time. The spacetime attractor instead instantiates multiple copies of a forward and inverse world model in the synapses between different neural populations (Methods). Recurrent dynamics can then arbitrate between many possible futures in parallel. Entorhinal grid cells also embed a world model in their connectivity (McNaughton et al., 2006), but they only encode a single location at a time (Vollan et al., 2026). Such networks can generate sequences, but the individual elements are represented one by one (Sompolinsky and Kanter, 1986; Kleinfeld, 1986; Widloski et al., 2025). The spacetime attractor suggests that circuit principles in prefrontal cortex resemble other cortical areas that use structural knowledge to infer features of the world. The major difference is that PFC instantaneously represents many points in time, which generalises known circuit principles to complex planning.

### STAs are flexible planners

Prefrontal cortex is particularly important for flexible behaviour in changing environments (Figure 1D; Burgess and Wu, 2013). To understand whether the spacetime attractor is a good model of planning in PFC, we therefore (i) study its behaviour and performance in such dynamic tasks, and (ii) characterise when and how it differs from existing models. We will see that the spacetime attractor is well-suited to dynamic ‘PFC-like’ problems, which other mechanistic models struggle to solve.

#### From static to dynamic tasks

We designed a set of four tasks that vary in how much the reward changes in space and time (Figure 4A). The tasks are all embedded in Euclidean space for simplicity of exposition, but the underlying principles generalise to other environments with known structure. Task 1 is simple navigation towards a static goal that remains constant across trials. In task 2, the static goal changes between trials. Task 3 is navigation to a goal that also moves within each trial, where each location is only rewarded when the goal is there. Task 4 generalises the idea of time-dependent goals to a non-binary reward landscape. Reward magnitudes are sampled independently between −1 and +1 for each location at each time point in each trial. The objective is to maximise cumulative reward over the trial, which requires balancing immediate reward with the potential for future reward (Figure 4A, right). This is reminiscent of the example from Figure 1D, where the restaurant and cinema are desirable at different times. To better understand how the STA solves these tasks, we compare it to two models commonly studied in systems neuroscience. The first is temporal difference (TD) learning, which has neural correlates in striatum (Sutton, 1988; Schultz et al., 1997). The second is the successor representation (SR), which has neural correlates in hippocampus (Dayan, 1993; Stachenfeld et al., 2017). These models assume fixed rewards across trials (TD) or within a trial (SR). Unlike the spacetime attractor, they do not generalise well when the reward changes rapidly.

**Figure 4.**
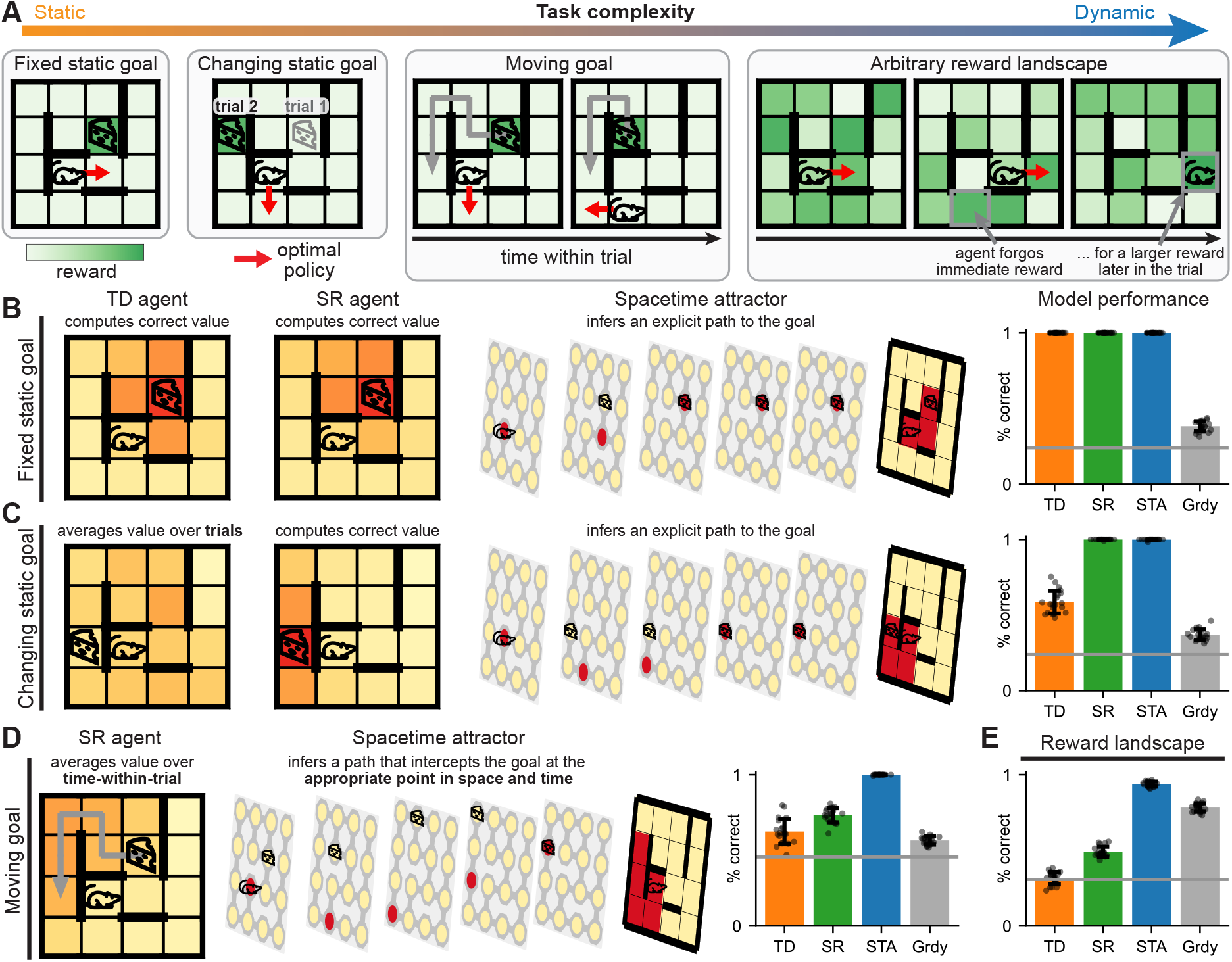
Model comparisons in static and dynamic tasks. **(A)** Example tasks with different degrees of dynamic reward. Colours indicate reward at each location (white to green), and red arrows indicate the optimal policy. In the ‘moving goal’ task, the agent intercepts a target that moves along a different trajectory in each trial (arrow). In the ‘reward landscape’ task, the reward is sampled independently in space and time. The agent has to maximise cumulative reward, and it can be optimal to forgo immediate reward for a later payoff. **(B)** Representations and performance in the static goal task with a fixed goal. Left: value functions learned by TD and SR agents. Centre: STA representation encoding a path through space and time. Right: performance of each model (see Figure S1 for STA performance in larger environments). The grey bar is a ‘greedy’ baseline that maximises immediate reward, and the horizontal line is a random baseline. **(C)** Task with a static goal that changes between trials. **(D)** Moving goal task. Left: the SR computes a value function that averages reward across time-within-trial. Middle: the STA takes into account the moving goal and computes a path that intercepts it. Right: performance of each agent. **(E)** Performance in the reward landscape task. All error bars indicate 1 standard deviation across 20 agents and environments (dots).

#### Simpler models can solve static tasks

TD learners gradually propagate value from rewarded locations to all other locations, and optimal policies are only learned when rewards are constant across both time and trials (Figure 4B-C, orange). The SR agent uses the environment structure and trial-specific reward function to compute values, and it can solve tasks where the goal changes between trials (Figure 4B-C, green). The STA can also solve these tasks, but it does so by inferring an entire spatial trajectory to the goal instead of computing a value function (Figure 4B-C, blue). Because the reward is constant throughout a trial, each sub-space receives the same reward input. However, the inferred representation differs across subspaces because it is also constrained to satisfy the current agent location and the transition structure of the world. As we will see, the inference of an explicit spacetime representation by the STA increases its flexibility compared to the amortised state values computed by TD and SR agents.

#### Only the STA solves dynamic tasks

While the SR agent can adapt to rewards that change across trials, the reward structure still needs to remain constant for the duration of the planning horizon. This is because the SR computes time-averaged occupancy and lacks fine-grained information about when the agent is where. When the reward changes within a trial, the SR averages values across timewithin-trial and fails to intercept the goal at the correct point in space and time (Figure 4D). The STA can solve the moving goal task because the input to each subspace is the reward at that specific moment in time. The representation is therefore biased towards coinciding with the goal in both space and time, and the network dynamics relax to a future trajectory that correctly intercepts the goal (Figure 4D). The advantage of the STA is exacerbated in the more dynamic reward landscape task, where very different locations can be rewarded at different times (Figure 4E). These results suggest that different algorithms could contribute differently to decision making. TD learning could drive rapid decisions in environments with stable rewards; the SR would be more flexible when rewards change on intermediate timescales; and the STA could enable adaptive behaviour in environments with rapid changes. Finally, sequential search would facilitate slower planning in novel environments and could be guided by partial plans from a spacetime attractor.

### STAs are efficient planners

We have now seen that spacetime attractors excel at dynamic planning problems. This required (i) a representation of the future, and (ii) structured weights that reflect the environment structure. However, it is not known whether this solution is optimal. In this section, we first train neural networks to infer optimal plans while imposing an explicit representation of the future. These networks learn to embed the environment structure in their recurrent weights. We then train fully unconstrained networks to generate optimal actions, and we show that they learn to infer explicit representations of the future using such structured connectivity.

#### Optimal weights for future representations

The representation and connectivity of the STA are inspired by theories of planning as an inference over future locations. However, the proposed weights may not be optimal for this computation. We therefore trained a recurrent network to perform inference over representations of the future, which were constrained to be factorised over time. The exact posterior for each future time step has a known functional form, which combines two factors that reflect constraints from earlier and later times in the plan. These factors can be approximated by recurrent inputs from earlier and later subspaces respectively (Derivation). However, we do not know the optimal form of the recurrent weights. We therefore optimised the weights numerically to maximise the similarity between (i) the exact posterior over future locations, and (ii) the fixed points of the network dynamics (Methods). The resulting connectivity resembled the handcrafted spacetime attractor (Figure S2). In particular, neurons representing consecutive time steps were connected according to the environment structure. Unlike the handcrafted STA, more distant populations were also weakly connected. This connectivity captured the higher-order structure of possible trajectories.

#### Optimal representation and connectivity

So far, we have directly imposed an explicit representation of the future. However, this representation requires many neurons. We investigated whether superior representations exist by training recurrent neural networks (RNNs) to solve dynamic planning problems efficiently (Mante et al., 2013; Stroud et al., 2023). We will see that regularised RNNs learn to solve these ‘PFC-like’ problems with both representations and connectivity that resemble a spacetime attractor, which suggests that it is an efficient solution. The RNNs were trained with supervised learning to generate optimal actions from inputs that indicated the current location and trial-specific rewards (Figure S3; Figure S4; Methods; Supplementary Note). The loss function only constrained the upcoming action and did not impose a representation of the distant future. To avoid biasing the RNN representation, reward information was only provided during initial ‘planning’ prior to the ‘execution phase’ of the task. The objective function also penalised large firing rates and network weights to encourage an efficient solution.

The defining features that allow a spacetime attractor to plan via inference are: (i) it has an explicit representation of the future; (ii) the connectivity embeds the environment structure; and (iii) attractor dynamics constrain the representation. RNNs trained on the reward landscape task exhibit all of these properties. These results are consistent with a prominent theory that PFC resembles a recurrent meta-learner, which learns connections over long timescales that rapidly adapt to new problems through recurrent dynamics (Wang et al., 2018). The STA similarly embeds stable structure in its weights and uses recurrent dynamics for fast planning with different rewards. We show that an STA is the mechanism implemented in the dynamics of meta-learners trained on challenging planning tasks.

#### RNNs learn explicit future representations

Spacetime attractors compute explicit representations of the future (Figure 5A-B). Critically, these representations generalise over trajectories. In other words, the representation in subspace *δ* depends only on the expected location in *δ* actions, and not on the rest of the trajectory (Figure 5A, right). To confirm that the RNN learned such a representation, we trained linear decoders to predict the future from its hidden state. To ensure generalisation across trajectories, we trained the decoders while holding out each ‘current location’ and computed the test accuracy only from those held out locations. The RNN had an explicit representation of the future during both planning and execution (Figure 5C-D; Figure S5). These representations were high-dimensional and explained substantial variance in the RNN activity (Figure S6). Consistent with experimental recordings from PFC (Chiang and Wallis, 2018), the STA and RNN both represented the future more strongly than the past (Figure 5D). In the spacetime attractor, the same subspace always encodes locations *δ* actions into the future. The future representation of the RNN also exhibited such ‘conveyor belt’ dynamics (Figure 5E). The neural representation of the RNN is thus consistent with a spacetime attractor.

**Figure 5.**
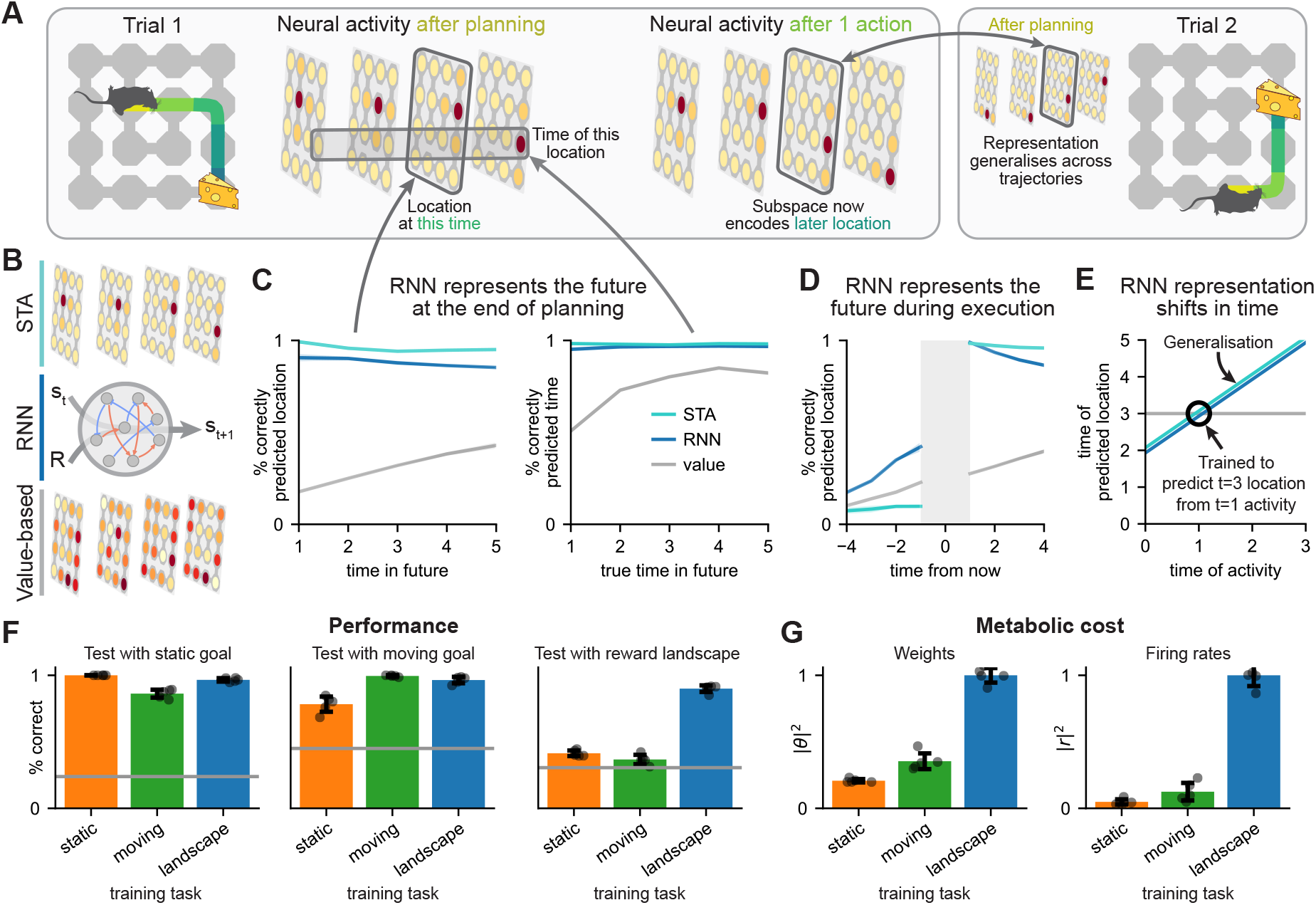
RNNs learn spacetime representations in the reward landscape task. **(A)** An STA infers the entire future during planning and represents the ‘future-to-go’ during execution (left). The representation in each subspace generalises across all trajectories passing through that location in spacetime (right). **(B)** We compare a spacetime attractor; an RNN trained on the reward landscape task; and an agent that computes an exact value function in space and time. The value-based agent computes an optimal policy from ‘neural activity’ containing (i) the value function, (ii) the agent location, and (iii) the time-within-trial (Methods). **(C)** Decoding accuracy at the end of planning for: left; agent location at each time in the future. Right; the time at which the agent will be at a given location, plotted as a function of the actual time the location was visited. Decoders were trained in crossvalidation across the current agent location (Methods). **(D)** Decoding accuracy during execution of location at each time in the past or future. **(E)** We trained a single decoder to predict location at time 3 from neural activity at time 1 (black circle). The same decoder predicted location at time *t* + 2 from neural activity at any other *t*, demonstrating ‘conveyor belt’ dynamics. **(F)** Performance in the static goal (left), moving goal (centre), and reward landscape (right) tasks for RNNs trained on either task (x-labels; colours). **(G)** Normalised parameter magnitudes of the three RNNs (left) and average firing rates in the static goal task (right). All error bars indicate 1 standard deviation across 5 RNNs (dots).

Intriguingly, RNNs trained on the simpler static and moving goal tasks did not reliably learn explicit future representations (Figure S7). Additionally, the RNN trained on the reward landscape task could solve these simpler tasks, while RNNs trained on the simpler tasks failed in the reward landscape task (Figure 5F). Alternative solutions therefore exist in the simpler tasks, and these solutions are favoured by a pressure to be energetically efficient (Figure 5G). We identify a tradeoff between generality and efficiency, where the PFC-like solution is general but needs more neurons and synapses, while simpler algorithms can solve simpler tasks more efficiently. Unlike value coding and related theories, this suggests a reason why PFC occupies such a large cortical territory.

#### RNNs learn a world model

In a spacetime attractor, subspaces that represent expected future locations are connected according to the environment structure. Testing this prediction in the RNN requires interpretation of the network weights, which is challenging because the functions of individual neurons are unknown. We therefore project the weights into a coordinate system, where the axes are orthogonal directions in neural state space that predict each point in spacetime (Methods). While this does not perfectly recover the true subspaces in the handcrafted STA, it is a good estimate. The projected RNN weights can therefore be interpreted as interactions between representations of different points in spacetime (Figure 6A).

**Figure 6.**
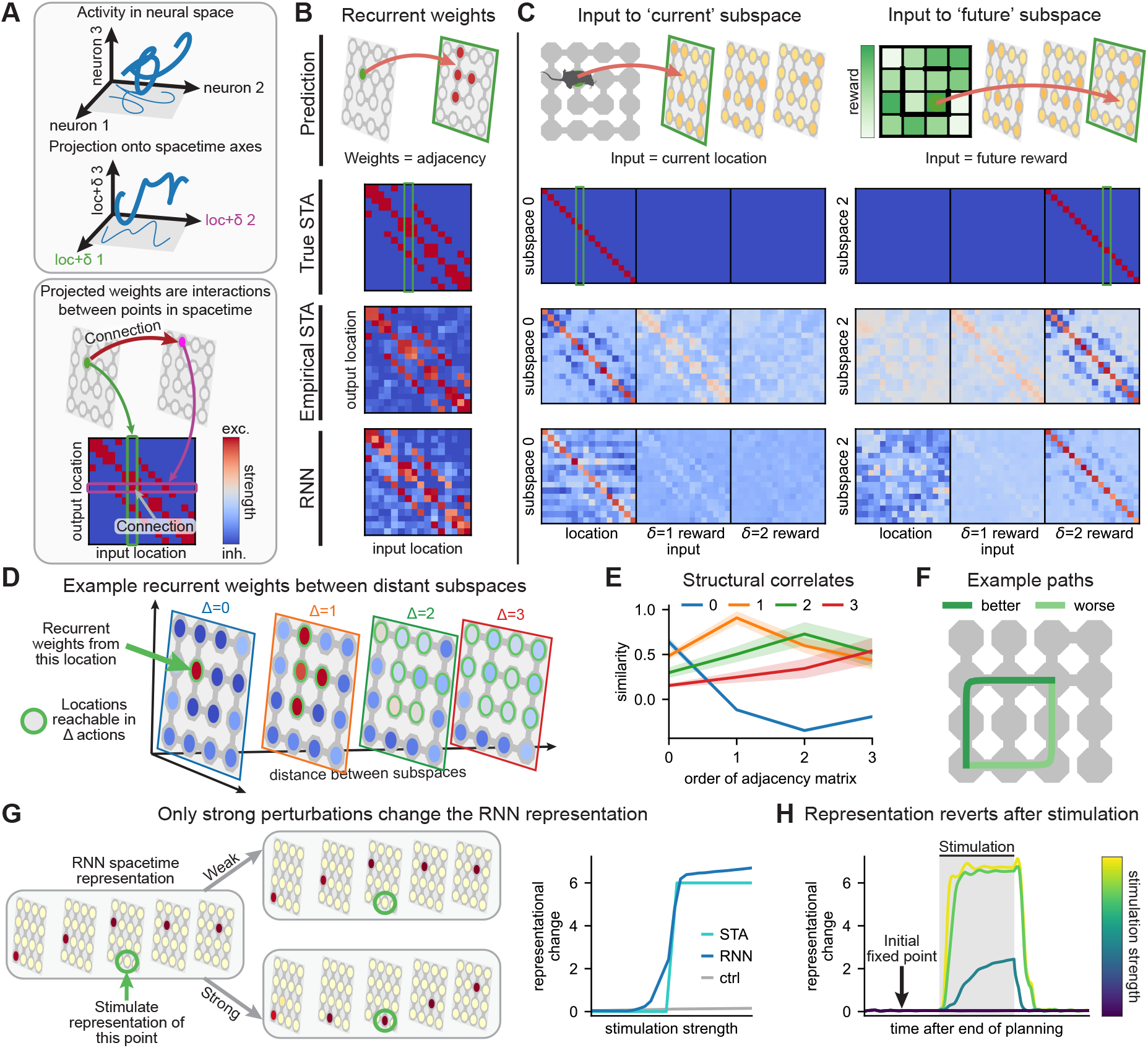
RNNs learn a world model. **(A)** The weights of RNNs trained on the reward landscape task are projected into an orthonormal coordinate system with axes that predict different points in spacetime (top). The projected weights are interactions between points in spacetime (bottom). **(B)** The average recurrent weights between subspaces separated by a single action resemble the environment adjacency matrix (bottom, ‘RNN’). ‘Empirical STA’ is the same analysis performed on approximate subspaces estimated from neural activity in the handcrafted STA. ‘True STA’ indicates weights between the ground truth future-coding subspaces in the handcrafted STA. Green box in ‘True STA’ indicates weights between (i) the single location denoted by a green circle in ‘Prediction’, and (ii) all locations in the subspace indicated by a green square. Input weights to the ‘current’ (*δ* = 0; left) and a ‘future’ (*δ* = 2; right) subspace. Consistent with an STA, the current subspace of the RNN receives location input, and future subspaces receive reward information. **(D)** Average recurrent weights between an example location (green arrow) and other locations at increasing time differences (Δ; planes). Locations in subspaces Δ apart are more strongly connected if they can be reached in Δ actions. Light green circles indicate these Δ^th^ order adjacency matrices (Δ = 0 corresponds to the identity). **(E)** Correlation between (i) the average connectivity between subspaces separated by Δ actions (lines; legend), and (ii) different order adjacency matrices (x-axis). Shading indicates standard deviation across 5 RNNs. **(F)** Example environment with a high-value path and a lower-value path. **(G)** Weak stimulation does not affect the RNN spacetime representation, but strong stimulation switches it to the lower-value path. **(H)** Magnitude of RNN representational change across time and stimulation strength (Methods).

Remarkably, the recurrent weights between adjacent subspaces closely resembled the adjacency matrix of the environment (Figure 6B; Figure S8; mean *±* std correlation of 0.91 *±* 0.07 vs. 0.72 *±* 0.06 for control environments). The RNN thus learned a world model explicitly in its recurrent weights. Consistent with a spacetime attractor, the *δ* = 0 subspace received input indicating the current agent location, and subspace *δ* received input indicating the reward in *δ* actions (Figure 6C). The RNN also learned weaker connections between more distant subspaces (Figure 6D-E). These connections reflected higher-order transition statistics, reminiscent of networks optimised explicitly to infer future representations (Figure S2). In summary, the RNN learned all of the structural components necessary to infer future actions by using a world model to integrate expected reward across time.

#### RNNs learn attractor dynamics

In a spacetime attractor, structured connections give rise to attractor dynamics with fixed points corresponding to explicit representations of desirable futures. To demonstrate similar dynamics in the trained RNN, we perturbed its neural activity at the end of planning. This resembles previous experiments in biological attractors (Inagaki et al., 2019; Kim et al., 2017; Vinograd et al., 2024). An attractor network should be robust to small perturbations, and it should be more sensitive to perturbations towards other plausible representations than perturbations along random directions in neural state space. We analysed the RNN in a setting with two ‘good’ paths that were close in value (Figure 6F). Like the handcrafted STA, the RNN initially converged to a stable representation of the better path, which persisted under weak perturbations (Figure 6G; left). Stronger perturbation of a single location on the alternate path caused a discrete change to a representation of this entire path, including in subspaces not directly perturbed (Figure 6G; right). This new representation remained after removing the perturbation in the handcrafted STA, suggesting that it is also a fixed point of the unperturbed dynamics (Figure S9). In contrast, the RNN representation relaxed back to the original fixed point after removing the perturbation (Figure 6H; Figure S9). The unperturbed RNN dynamics therefore only have a single stable fixed point, suggesting that the network may be less susceptible to local minima.

Together, our analyses show that RNNs trained on a dynamic planning task learn to approximate a spacetime attractor. This was also true across variations in model architecture (Methods; Figure S10; Figure S11). These results extend previous findings that explicit spacetime representations are optimal for sequence memory (Supplementary Note; Whittington et al., 2023; Dorrell et al., 2026; Wang et al., 2025). Additionally, RNNs with too few hidden units to learn a spacetime attractor failed to solve the task (Figure S12), suggesting that other solutions are not readily learned by gradient descent. The spacetime attractor is thus an efficient solution to dynamic planning problems, and it is consistent with a prominent theory that PFC can be understood as a recurrent meta-learner.

### STAs can generalise across environments

Hardwiring the transition structure of the environment into the synapses of a spacetime attractor seems prohibitive, because it might prevent the network from generalising across different structures (Figure 7A). Here, we show that is not the case. RNNs trained in environments with changing structure still learn spacetime attractors. The network dynamics are adapted to each environment through input-mediated gating of a generic scaffold. This is a simple mechanism for rapid adaptation that preserves the ability to plan under changing rewards.

**Figure 7.**
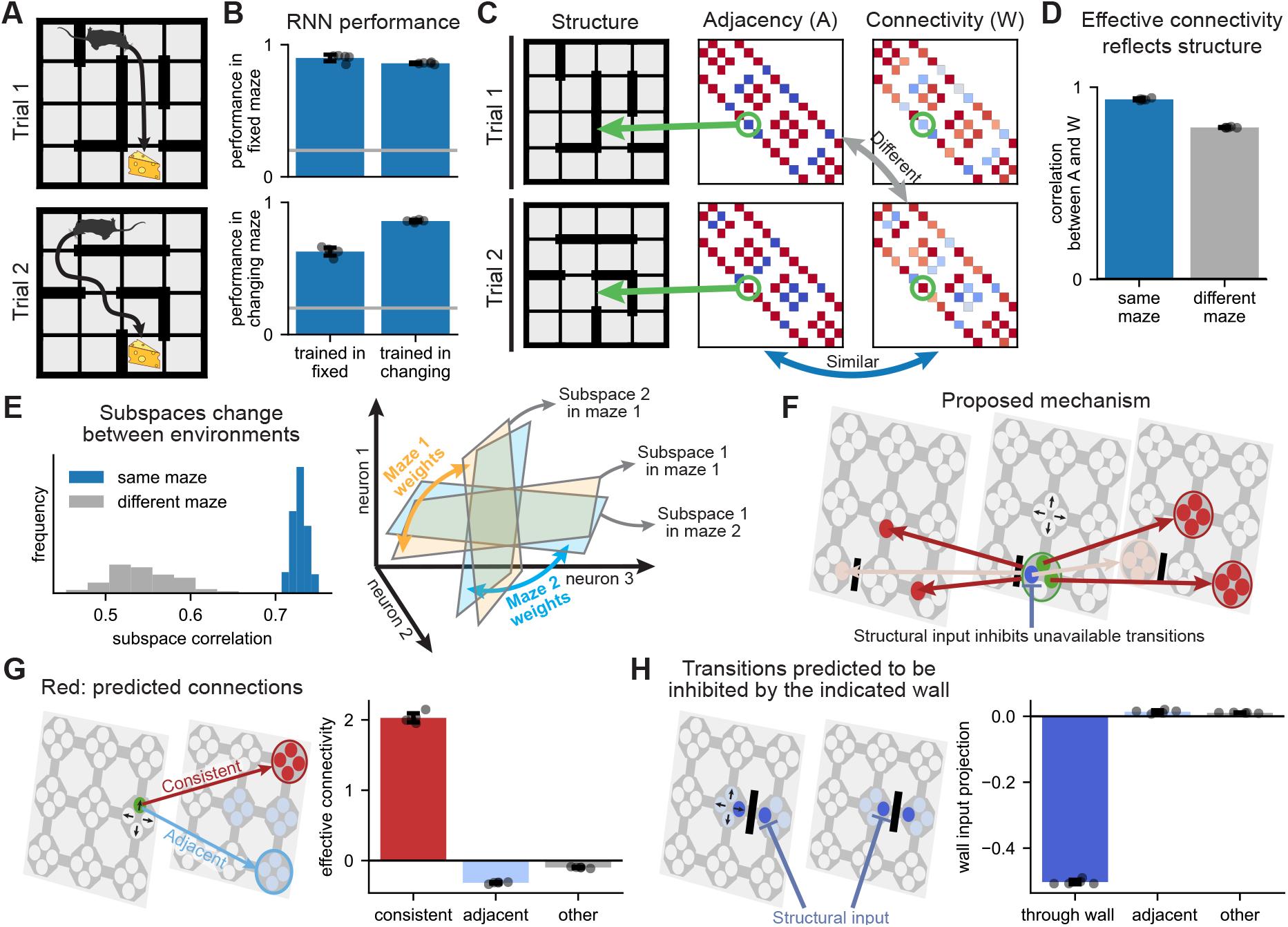
Spacetime attractors can adapt to changing structure. **(A)** Changing structure requires adaptation. **(B)** Performance of RNNs trained on the reward landscape task in a single maze or with a different structure on every trial, when evaluated in either a single maze (top) or across changing mazes (bottom). **(C)** Two example mazes (left) and their corresponding adjacency matrices (centre). The effective connectivity between adjacent subspaces in a single RNN (right) reflects the structure of each maze. **(D)** Average correlation between subspace connectivity in a maze and the structure of either the same (blue) or a random (grey) maze. **(E)** The spacetime representation is more similar across distinct trials from the same maze than trials from different mazes (left). By using slightly different subspaces in different environments, a network can match its connectivity to the environment structure (right; orange vs. blue). **(F)** Putative mechanism for structural generalisation. Instead of representing each future *location*, neurons encode expected future *transitions* (black arrows in example state). Each transition 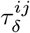 (green) excites other transitions that can follow 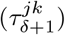 or precede 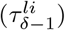 it (red arrows). Structural input to PFC inhibits transitions that are not available in a given environment (blue), preventing planning between states that would otherwise be connected (light red arrows). **(G)** Effective connectivity between directions in neural state space that encode ‘consistent’ consecutive future transitions (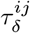 and 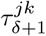; green to red), ‘adjacent’ transitions (green to blue), or any other transitions (green to grey). **(H)** Projection of the input from wall *w*_*ij*_ onto representations of future transitions *through* the wall (*τ*^*ij*^; dark blue), transitions to other adjacent states (*τ*^*ik*^; light blue), or any other transitions (grey). All error bars indicate 1 standard deviation across 5 RNNs (dots).

#### Representations adapt to the environment

To study planning in environments with changing structure, we trained RNNs on a variant of the reward landscape task, where the transition structure was different on every trial. The structure was provided to the RNN as a binary input that indicated whether otherwise adjacent states were separated by a wall (Methods). Importantly, the network had to adapt to each new environment through its dynamics while keeping the weights fixed across environments (Wang et al., 2018; Jensen et al., 2024). The generalising RNNs performed nearly as well as networks trained in a single environment (Figure 7B), raising the question of how generalisation is achieved. The neural representation resembled a spacetime attractor in any single environment (Figure S13B). However, the effective connectivity between subspaces changed between environments. Remarkably, the connectivity reflected the transition structure of whichever environment the agent was currently in (Figure 7C-D). This indicates a general solution that allows recurrent networks to use attractor dynamics for planning in any similar environment, without requiring synaptic plasticity.

#### A model of generalization

We can understand how the network adapts to the environment at both the ‘population level’ and at a mechanistic level. At a population level, the directions in neural state space that represent each point in spacetime were more similar on different trials from the same environment than across trials from different environments (Figure 7E, left). This change allows the network to align its representation with the appropriate components of the connectivity matrix (Figure 7E, right). How is this achieved mechanistically? We hypothesised that the network explicitly represents future *transitions* rather than future locations. In other words, there are directions in neural state space that represent ‘*in δ actions from now, move from state s*_*i*_ *to state s*_*j*_’, which we denote 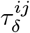. This generalises the vanilla spacetime attractor, which has a representation of future locations that is invariant to the subsequent state.

While it requires more neurons, explicitly representing future transitions allows the network to use input-mediated inhibition to adaptively ‘turn off’ transitions that are not available in any particular environment. The learned structure of the remaining transitions is then used for planning (Figure 7F). Importantly, the scaffold only needs to include transitions that are possible in at least one environment. This model makes three predictions that we verified: (i) the network explicitly represents future transitions (Figure S13C), (ii) the representations of future transitions are connected by a world model (Figure 7G), and (iii) input indicating a wall between *s*_*i*_ and *s*_*j*_ specifically inhibits representations of future transitions between those states (Figure 7H). These analyses show that the spacetime attractor is an efficient planning algorithm in structurally changing environments, and they suggest a general mechanism for adaptive behaviour.

## Discussion

We have taken inspiration from known cortical attractor dynamics and prefrontal sequence representations (Figure 1) to develop a new theory of planning. This spacetime attractor model uses different neural subspaces to represent the expected location of an agent at different times in the future. The subspaces are connected according to the structure of the environment, which allows the network to infer optimal future trajectories (Figure 2; Figure 3). The spacetime attractor solves problems with dynamic rewards, which PFC is important for and simpler algorithms struggle with (Figure 4). It solves these problems using internal representations that closely resemble prefrontal representations during sequence working memory, thereby unifying working memory and planning in PFC. RNNs trained to solve dynamic planning tasks learn to implement the spacetime attractor algorithm in their internal dynamics, suggesting that it is an efficient solution (Figure 5; Figure 6). Finally, spacetime attractors can generalise across environments with different transition structures through rapid gating of a general scaffold to reflect each particular environment (Figure 7).

### Experimental predictions

The spacetime attractor is inspired by data and makes precise predictions that can be tested in future experiments:

- After planning a behavioural sequence, different subspaces of PFC activity should represent distinct steps of the plan.
- Optogenetic activation of neurons in a future subspace should bias representations and behaviour towards the stimulated state at a delay corresponding to the stimulated subspace.
- The effective connectivity between subspaces should reflect the structure of the environment. This could be tested in noise correlations across neurons or by explicit holographic stimulation.
- Different patterns of neurons should represent the future in environments with different transition structures.
- The neurons active in each environment should be connected according to its structure. Inputs that mediate this gating could come through sensory cortex in observable environments or hippocampus when the structure is learned.

We believe that such hypothesis-driven experiments are critical to uncover mechanistic principles in prefrontal cortex. Although the reward landscape task is conceptually simple, RNNs trained on the task learn very high-dimensional representations (Figure S6; Figure S12). The underlying neural mechanism would be difficult to elucidate by studying low-dimensional projections of neural activity. However, the spacetime attractor provides a concise explanation that can be summarised in just a few equations (Methods). This enabled detailed understanding of both the representations, connectivity, and dynamics of the trained RNNs. We hope similar targeted analyses will uncover new mechanistic insights for flexible behaviour in biological circuits.

### Decision making across the brain

Our comparisons of different models across different tasks suggest complementary contributions of different brain regions to decision making. Striatum is thought to implement temporal difference learning (Schultz et al., 1997), which facilitates rapid responses in stable environments. Hippocampus embeds the structure of the world, but it is thought to either average representations over future actions in a successor representation (Stachenfeld et al., 2017) or represent one state at a time in a sequence (Widloski and Foster, 2022; Whittington et al., 2020; Jensen et al., 2024). This provides an efficient solution to problems with consistent spatial structure and rewards that change on intermediate timescales. A spacetime attractor in frontal cortex could facilitate adaptive behaviour in dynamic environments with a familiar structural scaffold that has been embedded in PFC. Finally, explicit search would facilitate slower planning in novel environments, and it could be guided by partial plans from a spacetime attractor. While spacetime attractors are particularly useful in dynamic environments, they can also solve simpler problems. It is therefore possible that animals have developed a spacetime attractor because they *sometimes* need it for planning, and then reuse it in simpler tasks. However, it is also plausible that many laboratory behaviours instead engage simpler algorithms in the basal ganglia or hippocampus. If that is the case, richer spacetime problems may be needed to query the representations used for planning in prefrontal cortex.

### Interactions with other planning algorithms

Planning via inference in a spacetime attractor differs from most models of planning in cognitive science. In particular, the spacetime attractor posits that planning can happen via *recognition*, where the sequence of steps needed to reach some desired state is directly inferred. In contrast, many studies of human planning focus on explicit search, where different paths are simulated and evaluated sequentially. We suggest that these two processes coexist, with spacetime attractor dynamics facilitating rapid planning-as-inference in familiar environments, where the structure has been embedded in prefrontal connections. The STA could also help focus explicit search towards putative high-value paths and evaluate the utility of different paths. This could happen through interactions with hippocampal activity sequences that have been proposed to facilitate planning (either replays or theta sequences; Foster, 2017; Jensen et al., 2024; Vollan et al., 2026). In particular, Jensen et al. (2024) suggest that PFC biases which sequences are generated by hippocampus. PFC would then update its representation to make high-reward sequences more likely in an iterative policy improvement process. Hippocampal replay has also been proposed to implement a ‘DYNA’ algorithm, where past experience is used to learn a model-free policy during rest or sleep (Mattar and Daw, 2018, 2026). This is complementary to both the spacetime attractor and explicit search. DYNA facilitates rapid decision making when the optimal policy remains stable across time, while decision-time planning allows adaptation to changing environments.

### Learning a spacetime attractor

In the hand-crafted spacetime attractor, we embedded copies of the environment adjacency matrix in the connections between every pair of consecutive subspaces. The RNN analyses show that such structure can be learned from repeated experience. However, learning copies of the same parameters independently for each pair of subspaces is inefficient. It would be more efficient to store a cache of experienced trajectories that can be used to learn all of the parameters. Hippocampal replay has been proposed to build cognitive maps via such interactions with prefrontal cortex (Bakermans et al., 2023; Ou et al., 2025). In particular, Ou et al. (2025) suggest that hippocampal replay consolidates structural information from hippocampus into cortex. Replay from hippocampus to PFC during sleep or rest could therefore provide a mechanism for learning all of the STA parameters from the same data.

### The state space of planning

We have assumed that planning happens at the level of individual locations in the environment. However, humans often plan in more abstract spaces, which improves efficiency by reducing the required planning depth. We also plan hierarchically by first computing an abstract plan that can be refined in increasing levels of detail (Eckstein and Collins, 2020). Multiple spacetime attractors operating at different levels of abstraction can be coupled to implement such a hierarchical planner (Figure S1D-E). One ‘abstract’ STA computes a high-level plan. A second STA treats the next abstract state as a goal and computes a detailed plan to get there. Stacking spacetime attractors in this way allows inference of plans that are exponentially long in the depth of the hierarchy, and therefore the number of neurons. This greatly improves efficiency over the linear scaling of a non-hierarchical STA.

### Environment interactions

We have focused on environments that evolve independently of the agent. The rewards and transition structure can therefore be estimated independently and provided as inputs to a spacetime attractor in PFC. Many problems involve environments that instead depend on the behaviour of an agent. In ‘key-door’ problems, moving through a locked door becomes possible only after picking up the key. We posit that such problems can be solved by structural gating of a learned scaffold, similar to the STA that generalises across environment structures (Figure 7). The structural gating would no longer be an external input, but instead a function of the spacetime representation itself. For example, activating a cell that represents ‘being at the key’ could disinhibit ‘transitioning through the door’ at a later time.

Another important example of environment interactions is social behaviours, where agents can change their plans in response to each other. A system of two spacetime attractors could simulate the joint dynamics of two such agents. The first STA ‘A’ infers future behaviour for agent A. The reward input would be the output of a different STA ‘B’ that predicts the behaviour of another agent. The reward for STA B would itself be an output of STA A, which couples the predicted behaviour of the two agents. The combined system relaxes to a fixed point, where the behaviour of A is optimal given the predicted behaviour of B and vice versa – a putative neural implementation of ‘theory of mind’.

### Outlook

We have developed a new theoretical framework for planning in prefrontal cortex. It builds on recently discovered prefrontal working memory representations and known attractor dynamics in other neural circuits. The spacetime attractor extends these principles to prospective behaviours in complex and changing environments – a setting that has previously eluded mechanistic circuit models. Existing data does not allow us to test these ideas explicitly. Instead, we have provided a series of concrete predictions for future experiments. We hope this will inspire new work in both experimental and computational neuroscience.

## Author contributions

KTJ and TEJB developed the conceptual framework with input from TA, PD, MSM, SR, AB, and MS. KTJ performed the simulations and analyses. KTJ made the figures with help from MSM, SR, and AB. KTJ and TEJB wrote the manuscript with input from all authors.

## Code availability

Code to reproduce results and run example models is available at github.com/KrisJensen/pysta.

## Acknowledgments

We are grateful to Diksha Gupta, Mohamady El-Gaby, Will Dorrell, and Tom Mrsic-Flogel for helpful feedback on the manuscript. We thank Qianqian Feng for discussions relating to planning-as-inference. This work was supported by a Wellcome Principal Research Fellowship (219525/Z/19/Z; TEJB, KTJ, AB), a Royal Society Faraday Discovery Fellowship (FDF\S2\251056; TEJB, KTJ, AB), the Gatsby Initiative for Brain Development and Psychiatry (GAT3955; TEJB), the Jean Francois and Marie-Laure de Clermont Tonerre Foundation (TEJB), a Wellcome Trust Career Development Award (225926/Z/22/Z; TA), the Oxford Clarendon Fund (PD), the Human Frontier Science Program (LT0040/2024-L; SR), EMBO (ALTF 651-2023; SR), the Gatsby Charitable Foundation (GAT4058; MS), the Simons Foundation (SCGB543039; MS), and a FYSSEN postdoctoral study grant (MSM). KTJ, TEJB, MSM, and SR were also supported by the Sainsbury Wellcome Centre’s core provided by Wellcome (219627/Z/19/Z) and the Gatsby Charitable Foundation (GAT3755).

## Methods

### The spacetime attractor

#### Problem specification

We focus on the problem of planning, which can be defined as computing a sequence of actions that maximises expected future reward. To solve this problem, it is necessary to know (i) the current state *s*_*t*_ of the agent; (ii) the transition structure of the environment; and (iii) which states are likely to be rewarding in the future. We assume that the transition structure remains constant throughout each trial and use ***A*** ∈ ℝ^*N×N*^ to denote the *adjacency matrix* of the environment. Here, *N* is the number of states, and ***A*** has elements *A*_*ij*_ = 1 if it is possible to move from state *s*_*t*_ = *j* to *s*_*t*+1_ = *i*, and *A*_*ij*_ = 0 otherwise. We allow the reward function ***R*** ∈ ℝ^*T×N*^ to vary over the course of a trial, such that *R*_*δi*_ indicates the reward associated with visiting state *s*_*t*+*δ*_ = *i* in *δ* actions, and *T* is the fixed length of the planning horizon. The challenge of estimating future rewards from past data is beyond the scope of this paper, and we simply assume access to a ground-truth reward function. Instead, we show how a recurrent neural network can embed ***A*** in its weights and take *s*_*t*_ and ***R*** as inputs to compute a trajectory *{s*_*t*_, *s*_*t*+1_, …, *s*_*t*+*T*_ *}* with high cumulative reward 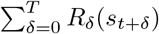.

#### Network dynamics

The spacetime attractor is a recurrent neural network with an exponential nonlinearity and block-wise normalisation. We define *z*_*δi*_ as the ‘potential’ and *r*_*δi*_ as the ‘firing rate’ of a neuron that represents a plan to ‘be at location *s*_*t*+*δ*_ = *i* in *δ* actions’. Each iteration of network dynamics then involves three steps:

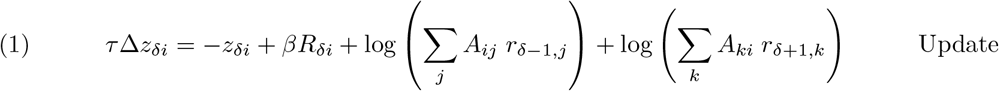

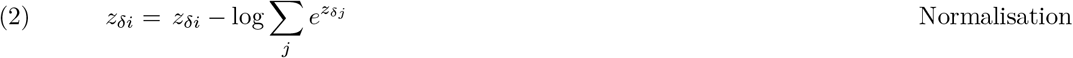

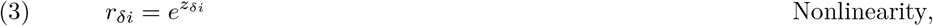

Here, *τ* = 50 network iterations is the ‘time constant’ governing the exponential decay of the neural potential in the absence of inputs. *A*_*ij*_ are weights corresponding to the adjacency matrix, and *R*_*δi*_ is an input that reflects the reward available at location *s*_*t*+*δ*_ = *i* in *δ* actions. To anchor the plan at the current agent location, we set *R*_0*i*_ = 20 if *s*_*t*_ = *i* and 0 otherwise. *β* is a temperature parameter that determines how strongly the agent values reward, and we set it to *β* = 9 unless otherwise noted. We take the policy of the agent to be greedy in the representation in subspace *δ* = 1 of locations accessible from the current state: *a* = argmax_*i* (*s*)_ [*r*_1*i*_]. The action *a* specifies which location to move to next. We used a maximum planning horizon of *T* = 6 for all analyses in the main text, a planning horizon of *T* = 16 in Figure S1B, and a planning horizon of *T* = 32 in Figure S1C. The normalisation in line (2) can be implemented through a subspace-specific inhibitory neuron with a logarithmic nonlinearity 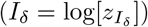, which receives input from all excitatory neurons in the subspace 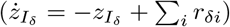 and provides reciprocal inhibition. In practice, we normalised analytically for simplicity of implementation.

The fixed points of the STA dynamics have the form

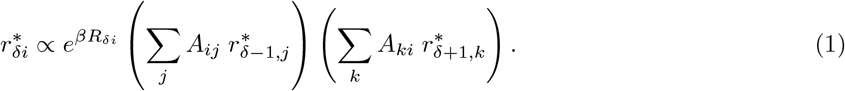

At convergence, the representation is therefore biased towards high-reward states (first term), while also being constrained to represent continuous trajectories (second and third terms).

#### Relationship to planning-as-inference

Here we briefly outline how the STA dynamics are inspired by previous models of ‘planning-as-inference’. Following Botvinick and Toussaint (2012), we formulate planning as an inference problem in a Markovian probabilistic graphical model. We assume a uniform prior over possible actions, such that 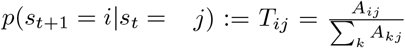. In line with Solway and Botvinick (2012) and Levine (2018), we define observations *o* ∈ [0, 1] as binary ‘optimality variables’ with emission probabilities

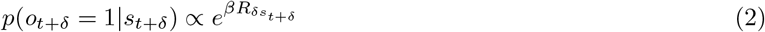

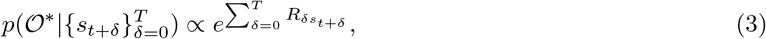

where we define 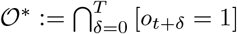 as the condition that *o*_*t*+*δ*_ = 1 for all *δ*. States with high reward are therefore more likely to have *o* = 1 than states with low reward, and the probability that *all* observations take a value of 1 is monotonic in the reward associated with the entire future trajectory. We can now *condition* on *O*^∗^ to compute a posterior distribution over future states:

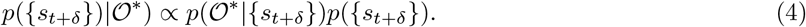

The posterior marginals can be computed with the forward-backward algorithm, but that requires three separate neural populations for each time in the plan to represent forward messages, backward messages, and their product. Instead, we define a *factorised* distribution *Q*(*s*_*t*+*δ*_) = Π_*δ*_*q*_*δ*_(*s*_*t*+*δ*_) and optimise it to resemble the true posterior as closely as possible. We assume that each component of this distribution is encoded in the firing rates ***r***_*δ*_ ∈ ℝ^*N*^ of a single population of neurons, such that *q*_*δ*_(*s*_*t*+*δ*_ = *i*) := *r*_*δi*_.

It is common to factorise the true posterior distribution over trajectories as:

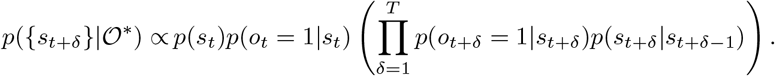

However, we can instead ‘split’ the transition terms across consecutive factors in a symmetrised factorisation:

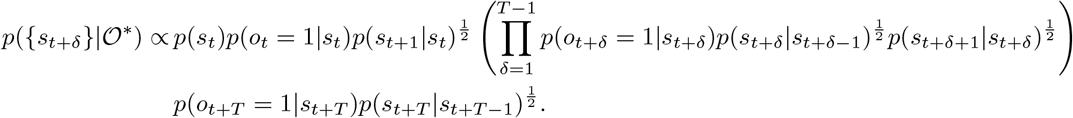

We can therefore define a ‘factor’ 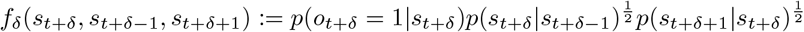 for each time in the plan. While the approximate *q*_*δ*_ only depends on *s*_*t*+*δ*_, *f*_*δ*_ also depends on *s*_*t*+*δ*−1_ and *s*_*t*+*δ*+1_. We want our approximate posterior to resemble the product of true factors as closely as possible:

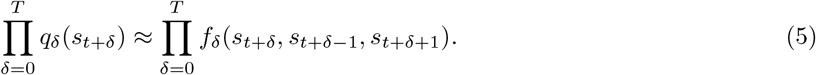

Loosely inspired by expectation propagation Minka (2013), we choose to *locally* minimise the discrepancy between the true and approximate posteriors. In other words, we define an objective function for component *δ* as the KL divergence between (i) the factor *f*_*δ*_(*s*_*t*+*δ*_, *s*_*t*+*δ*−1_, *s*_*t*+*δ*+1_), while approximating the distributions over *s*_*t*+*δ*_*′*_≠*δ*_ by *q*_*δ*_*′* (*s*_*t*+*δ*_*′*); and (ii) the distribution *Q*({*s*_*t*+*δ*_*}*) implied by the firing rates {***r***_*δ*_*}*. We also introduce a normalisation constant *Z*_*δ*_ to the first distribution. This gives rise to a loss function of the form

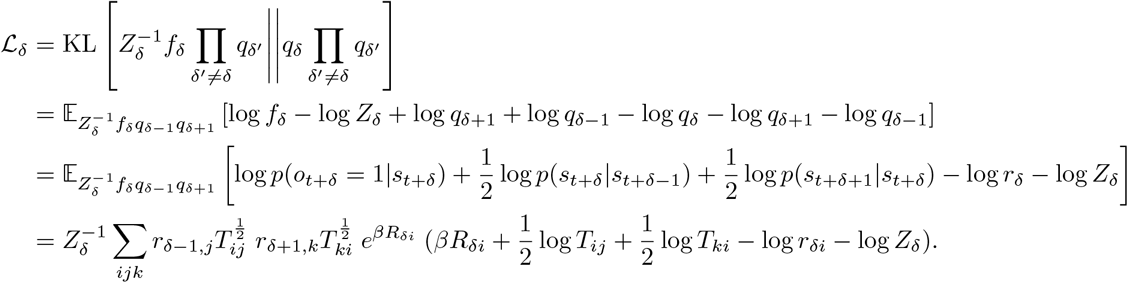

Here 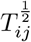 indicates an element-wise square root of each term in the transition matrix. We introduce a Lagrange multiplier *λ* for normalisation and take derivatives to minimise this objective:

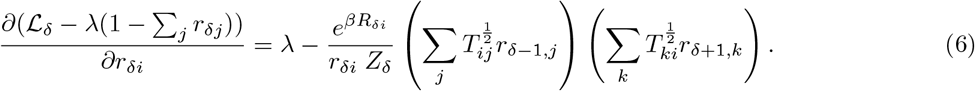

This local optimisation of a factorised approximate posterior leads to the ‘mode-seeking’ dynamics seen in Figure 3B. The fixed points have the form

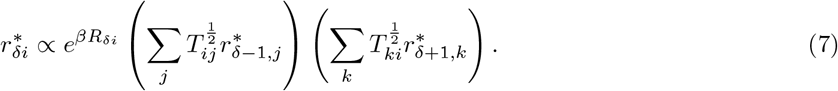

We now recall the STA fixed points:

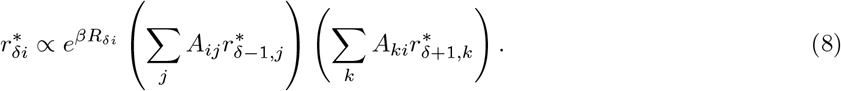

The difference between the two is that the STA uses the adjacency matrix ***A*** in place of the element-wise square root of the transition matrix, 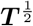. The STA therefore has the form of a dynamical system with fixed points that locally approximate the posterior marginals over trajectories, conditioned on ‘optimality’ in the planning-as-inference formalism. It also has very similar dynamics to another neural implementation of planning-as-inference by Donnarumma et al. (2025). However, the local approximation with marginals may be quite poor as it neglects parts of the transition matrix. For this reason, Figure S2 and Derivation directly optimise the dynamics of a recurrent network for inference.

#### Practical considerations

Here we note a few deviations from a simple interpretation of the STA dynamics as approximate inference.

First, we use the adjacency matrix instead of the transition matrix or its square root, which differs from a probabilistic interpretation. We do this because the true posterior distribution favours trajectories that pass through low-degree states, which have higher probability under the prior. In contrast, the recurrent weights between RNN subspaces representing consecutive future actions in Figure 6 were generally more similar to the adjacency matrix than the transition matrix. One way to interpret the adjacency matrix is as an action-optimised transition matrix, *A*_*ij*_ = max_*a*_ *p*(*s*_*t*+1_ = *i*|*s*_*t*_ = *j, a*_*t*_ = *a*). This generalises the idea to stochastic environments, but it does not optimise a well-defined objective to the best of our knowledge. All results in the paper hold qualitatively when using 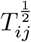 in place of *A*_*ij*_ for an implementation that is more faithful to planning-as-inference.

Second, we add a small amount of noise *η*_*ij*_ ~ *U* (−0.025, −0.015) to each element of the weight matrix instead of using the exact adjacency matrix. Noise is added because it is unreasonable to expect biological systems to exactly recapitulate the transition structure of the environment. It is restricted to be negative because the STA can be susceptible to discontinuous plans if noisy weights imply a finite probability of transitioning between two distant states (Supplementary Note). We similarly added white noise *ξ* ~ *N* (0, *σ* = 0.1) to the potential of all neurons at each iteration of the network dynamics.

Third, we threshold both the neural potentials (*z*_*δi*_), the forward messages (log ∑_*j*_ *z*_*δi j*_ *A*_*ij*_ *r*_*δ*−1,*j*_), and the backward messages (log ∑_*k*_ *A*_*ki*_ *r*_*δ*+1,*k*_) at a minimum value of *ϵ* = −100 for numerical stability.

Finally, the STA representation updates automatically when the agent moves, since both the location and reward inputs change. However, an additional ‘feedforward component’ in the form of the identity matrix was added to the connectivity between subspaces *δ* and *δ* − 1 during the first 100 network iterations (2 time constants) after every action. This is inspired by the ‘update neurons’ of the fruit fly head direction circuit (Turner-Evans et al., 2017) and stabilises the ‘conveyor belt’ dynamics, but it is not necessary for any of the main results in the paper (Supplementary Note). We ran the STA dynamics for 400 iterations before each environment update to ensure convergence to a stable representation.

### Optimised approximate planning-as-inference

In the handcrafted STA, we imposed both a representation and connectivity inspired by planning-as-inference. However, our derivation of approximate inference absorbed some of the transition terms into the approximate marginals. The handcrafted connectivity may therefore not be optimal for inference of future representations. Here we describe our approach to learning the weights of a recurrent network that optimally approximates planning-as-inference with a factorised distribution (Figure S2). The network dynamics are given by

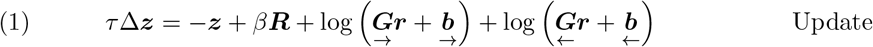

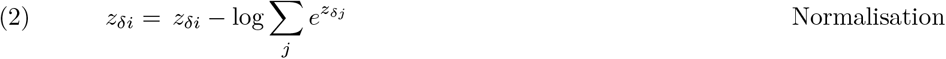

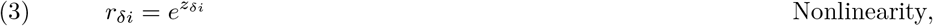

where 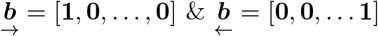 represent our uniform boundary conditions. Here we use ***R, r*** ∈ ℝ^*NT×*1^ to denote the vectorised versions of the reward function and neural activity across all times and locations. 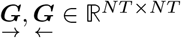 are constrained to be lower and upper block triangular respectively, and the STA dynamics are recovered if we set their off-diagonal blocks to the adjacency matrix.

The fixed points of these dynamics are given by

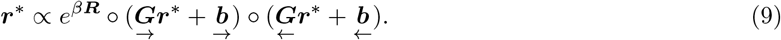

We optimised these fixed points to minimise the KL divergence between (i) the true marginal posterior *p*_*δi*_ = *p*(*s*_*t*+*δ*_ = *i*|*O*^∗^) at each delay *δ*, and (ii) the approximate posterior 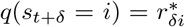 implied by the firing rates ***r***^∗^. The gradient of this objective with respect to the parameters is given by

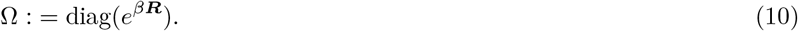

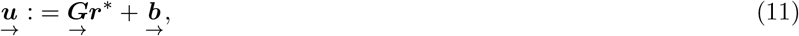

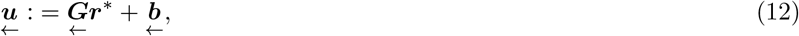

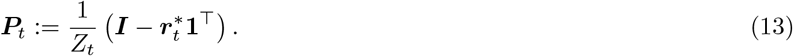

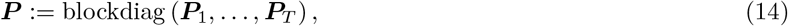

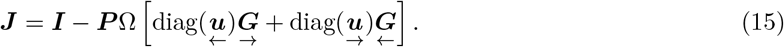

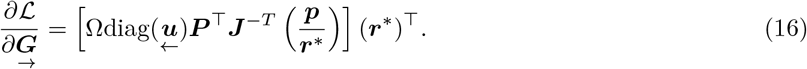

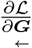 is given by the symmetric expression (see Derivation for details).

For each parameter update, we ran a batch of 200 trials from the reward landscape task with *τ* = 10 for a maximum of 500 network iterations or until convergence (max_batch,*δ,i*_ [abs 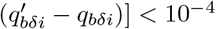). We then calculated the average gradient for 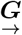 and 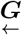 across the batch and used ADAM to optimise the parameters. After each parameter update, we projected 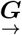 and 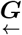 back onto the set of positive lower and upper block triangular matrices.

### Tasks

We used three different classes of tasks to train and compare models in this work – the ‘static goal’ task, the ‘moving goal’ task, and the ‘reward landscape’ task. All tasks required agents to navigate mazes on a 4 *×* 4 grid with *N* = 16 states (except the analyses of larger environments in Figure S1). Walls prevented transitions between some pairs of otherwise adjacent states, and the wall configurations were sampled as described by Jensen et al. (2024). For most analyses, the wall configuration remained fixed across all trials, and only the reward function changed. For the analyses in Figure 7, the wall configuration changed across trials, and the agent had to adapt to the new transition structure through its recurrent network dynamics. In all tasks, the agent location was sampled randomly at the beginning of each trial. At the beginning of each trial, the agent location and reward function were ‘frozen’ for 5-7 (randomly sampled on each trial) iterations of the environment. This constituted an initial ‘planning’ phase that was followed by an ‘execution’ phase, where (i) the agent took actions that changed its location, and (ii) the reward function updated as described in more detail below. For all analyses, the full reward function was provided to the agent during the initial planning period. For the handcrafted STA and the RNN in Figure S10, the input provided reward information that was relative in time during execution – it indicated what the reward would be at a given location *in δ steps* rather than *at time t*. The reward input was zero for all *δ* that corresponded to time points beyond the end of the trial. For all other RNN analyses, the reward input was set to zero during execution (Figure S3).

#### Moving goal task

In this task, a full goal ‘trajectory’ was sampled on each trial. The goal trajectory was sampled as a restricted random walk that could only turn around if it reached a dead end. The start location of the agent was restricted to not coincide with the start location of the goal. The trial terminated when the agent was at the same location as the goal at a given moment in time, or after a maximum of *T* = 6 actions. The reward input to the agent, ***R*** ∈ ℝ^*T×N*^, was a matrix indicating the location of the goal at every point in the future. The reward input consisted of ‘+0.6’ for the goal location and ‘-0.6’ for all other locations at each moment in time. In other words, *R*_*δi*_ was ‘+0.6’ if the goal would be at location *s*_*t*+*δ*_ = *i* after *δ* actions, and −0.6 otherwise.

#### Static goal task

This task was identical to the moving goal task, except that the goal remained stationary for the duration of each trial. The static goal task was implemented in two different variants – one where the goal remained fixed across all trials, and one where the goal was resampled at random in every new trial. The handcrafted models were evaluated in both of these tasks, while the RNNs were only trained with a goal that changed between trials.

#### Reward landscape task

In this task, every element *R*_*ti*_ of the reward function ***R*** ∈ ℝ^*T×N*^ was sampled uniformly and independently between −1 and +1 at the beginning of each trial. The complete future reward structure was provided as an input to the agent as described above. Every trial finished when the agent had taken *T* = 6 actions.

#### Quantification of performance

For most performance comparisons, we computed the probability of choosing the optimal first action in each trial. We define the optimal first action as the first action of the trajectory that maximises cumulative reward over the entire trial. We use this metric rather than the cumulative reward for two reasons. (i) We are interested in the process of planning, whereby an agent balances immediate and long-term reward. This is most challenging for early actions, while the greedy policy is optimal for the last action. (ii) The probability of choosing an optimal action is readily interpretable as a number between 0 and 1. All results were qualitatively similar if we instead used the probability of choosing an optimal action averaged over the entire trial as a performance metric, or if we used the average empirical reward.

For the analyses in Figure S1, we analysed whether the entire trajectory implied by the STA representation was correct. A representation was considered correct if the location with the highest firing rate in each subspace (i) was adjacent to the previous location, (ii) was closer to the goal than the previous location, and (iii) had a firing rate of at least 0.3.

### Reference models

Here we provide details of the temporal difference and successor representation baselines that we compare the spacetime attractor to. For further details, we refer to Jensen (2023). Common to both of these frameworks is that they explicitly estimate the ‘value function’ under some policy *π*:

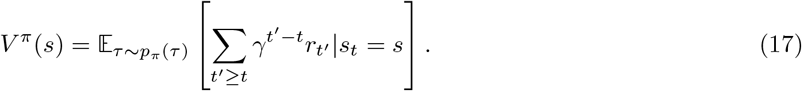

Here, 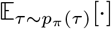 indicates an expectation taken over trajectories *τ* resulting from the agent following *π*. The ‘TD’ and ‘SR’ heatmaps in Figure 4 show the computed value functions. In both cases, we take the policy of the agent to be greedy in the value function evaluated at all locations accessible from the current state.

#### Temporal difference learning

We implement vanilla temporal difference learning, which computes a value function by iteratively applying the update

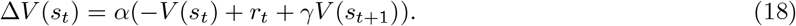

We set the temporal discount factor to *γ* = 1, since we are interested in maximising the non-discounted cumulative reward. We allowed the TD agent to interact with the environment for 4,000 trials with a learning rate of *α* = 0.05 before analysing its performance.

#### Successor representation

In the successor representation, the value function is decomposed as

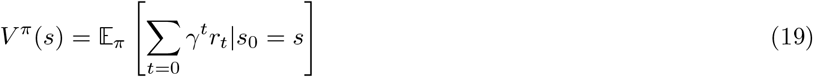

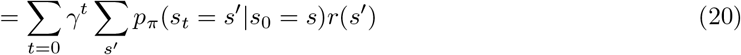

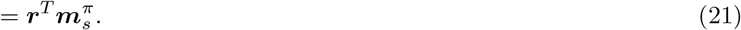

Here, ***r*** is a vector of the average reward associated with each state, and 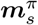 is a vector of the expected discounted future occupancy of state *s*^*′*^ if the agent starts in state *s* and follows policy *π*:

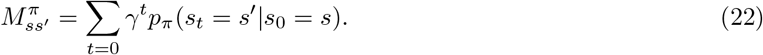

The full matrix ***M*** ^*π*^, constructed from stacking the 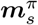 corresponding to all states *s*, is denoted the ‘successor matrix’, and it allows us to write down a vector of expected rewards from all states as

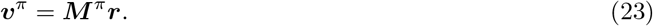

***M*** ^*π*^ can be learned using a TD-like algorithm as above. However, since the STA algorithm also has explicit access to the transition structure of the environment, we simply computed the exact successor matrix as the geometric series ***M*** ^*π*^ = ***I*** + *γ****T*** ^*π*^ + *γ*^2^(***T*** ^*π*^)^2^ + … = (***I*** − *γ****T*** ^*π*^)^−1^ with *γ* = 0.95. When computing the successor matrix, we take *π* to be the diffusion policy, and ***T*** ^*π*^ is therefore the diffusion matrix. This is similar to how the adjacency matrix serves as a global structural prior in the STA.

#### Spacetime value agent

In Figure 5, we compare representations to an agent that computes a value function in a state space consisting of space *and* time. This agent uses dynamic programming to compute the value of being at location *s* at time *t* under an optimal policy for every combination of *s* and *t*:

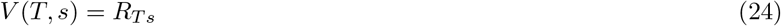

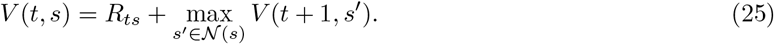

At time *t*, the optimal policy is greedy in the value of accessible locations at time *t* + 1. For the decoding analyses in Figure 5, we took the ‘neural representation’ to be the concatenation of (i) the flattened value function ***v***_*f*_ ∈ ℝ^*TN*^, (ii) a one-hot representation of the current location, and (iii) a one-hot representation of the current time-within-trial. These three quantities are sufficient to compute an optimal policy.

### Recurrent neural networks

In this section, we explain how recurrent neural networks were trained and analysed. All networks were fully connected with *N*_rec_ = 800 hidden units (except Figure S12) and ReLU nonlinearities. Each iteration of network dynamics was given by

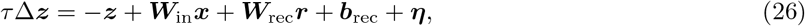

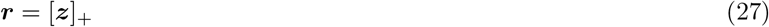

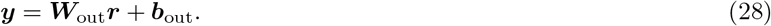

Here, 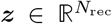 are the network potentials, 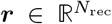 are the firing rates, 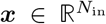 are the inputs, 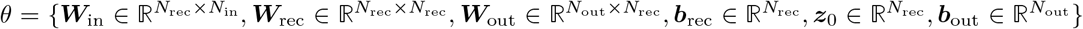 are the network parameters, and 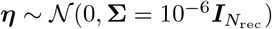 is Gaussian noise. All simulations used a time constant of *τ* = 5. The output policy 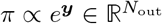 was defined in global allocentric coordinates as the desired next state (*N*_out_ = *N*). In Figure S11, we also analyse a network that produced a policy in ‘local’ coordinates that indicated the desired movement direction (*N*_out_ = 5). The input consisted of (i) a one-hot representation ***x***_loc_ ∈ ℝ^*N*^ of the current location in the environment; (ii) the reward ***x***_rew_ ∈ ℝ^*NT*^ as described for the tasks above; and (iii) a binary representation ***x***_*w*_ ∈ ℝ^2*N*^ of the location of all walls (Jensen et al., 2024). The elements of ***x***_*w*_ indicated for each of the *N* locations whether there was a wall (i) to the right of, and (ii) above it. In each trial, a subset of these were present, and a subset were absent. All ‘present’ walls were assigned a value of +1 in ***x***_*w*_, and all absent walls were assigned a value of 0. The total input dimensionality is therefore *N*_in_ = *N* (1 + 2 + *T*). Gaussian noise with a standard deviation of *σ* = 10^−3^ was added to the inputs before passing them to the RNN. A schematic illustration of all RNN inputs and outputs is provided in Figure S3.

For all analyses in the main text, the RNN performed 10 network iterations per environment iteration (action). Reward input was only provided during the planning phase and set to 0 during the execution phase (Figure S3). This was done to avoid biasing the execution-time representation towards a ‘relative’ subspace representation by providing reward input in that format (Supplementary Note). For comparison with this ‘working memory’ RNN, we also trained an RNN with reward input during both the planning and execution phases, which learned qualitatively similar representations and dynamics (Figure S10). For this network, the reward input was given relative to the current time as in the handcrafted STA, and the number of network iterations was randomly sampled between 9 and 11 before each environment iteration.

We optimised all network parameters *θ* to minimise the loss function

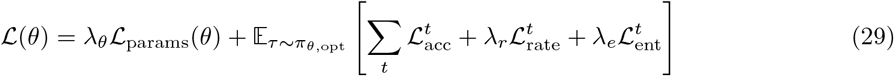

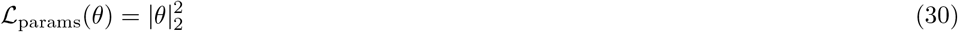

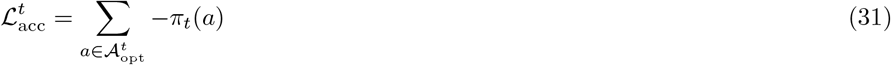

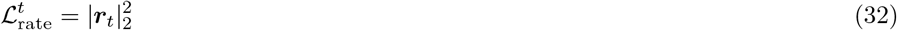

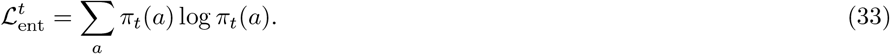

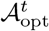 is the set of optimal actions at time *t*, and 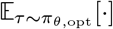 indicates an expectation over trajectories induced by the policy of the agent, renormalised over optimal actions. That is, we consider an imitation learning setting where ties between equally optimal actions are broken according to the actual policy of the agent. We used *λ*_*θ*_ = 2 *×* 10^−7^, *λ*_*r*_ = 10^−5^, and *λ*_*e*_ = 10^−4^ for all analyses. The RNNs were trained using Adam (Kingma and Ba, 2015) with a learning rate of 3 *×* 10^−4^ for 200,000 batches of 200 trials (250,000 batches for the RNNs trained in changing mazes). All results are reported as mean and standard deviation across 5 separate RNNs that were trained from different random seeds and with different environment transition structures.

### Analyses

In this section, we describe the analyses used to compare different handcrafted models and trained RNNs.

#### Fixed point analyses

For each analysis in Figure 3, we ran the STA for 1200 network iterations (24 time constants) with recurrent noise *ξ* ~ *N* (0, *σ* = 2.0) and no structural noise added to the weights. To quantify the frequency of convergence to different fixed points, we repeated each simulation 1000 times from a uniform initial representation. A simulation was considered to converge to a particular reference representation if the appropriate location in each subspace had a final firing rate of at least 0.9. For the analysis in Figure 3F (bottom), we set the initial neural potential in subspace *δ* = 2 to *z*_2*i*_ = 0 for the bottom right location and *z*_2*i*_ = − 100 for all other locations. The remaining subspaces were initialised to uniform representations. For the simulation in Figure 3E, we initialised the neural potentials to the final state from Figure 3B, set all reward inputs to zero, and ran the network dynamics for 1200 iterations.

#### Decoding analyses

To decode future locations from neural activity (Figure 5C-D), we used scikit-learn (Pedregosa et al., 2011) to train L2-regularised logistic regression models that predicted location at time *t*_*L*_ from neural activity at time *t*_*N*_ with an inverse regularisation strength of *C* = 1.0. We performed this analysis in crossvalidation across locations at time *t*_*N*_. In other words, we trained a decoder on data where the agent was in any state *s*_*t*_ ≠ *s* to predict all locations at time *t*_*L*_, and we then tested this decoder in trials where the agent was in state *s* at time *t*_*N*_. We repeated this analysis across all test locations *s* and averaged performance across the resulting 16 folds. We did this to test whether the RNN had a generalisable representation of future location, rather than an encoding of e.g. current location and neighboring values. In Figure 5C (left), we plot the performance of decoders trained on *t*_*N*_ = −1 to predict location at any *t*_*L*_ ≥ 1. In Figure 5D, we trained decoders on every pair of *t*_*N*_ ∈ [0, 5] and *t*_*L*_ ∈ [0, 5]. We then averaged the performance across every ‘delay’ *t*_*L*_ − *t*_*N*_.

To investigate the generalisation of decoders in Figure 5E, we trained a single decoder using neural activity at time *t*_*N*_ = 1 to predict location at time *t*_*L*_ = 3. We then applied the same decoder to neural activity at all times 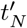 and quantified how well it predicted location at all times 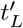. This analysis was again performed in crossvalidation. A separate decoder was trained while holding out each location *s*_1_ = *s*, and then tested only on trials where 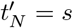. Performance was averaged across all held-out locations and across 5 independently trained RNNs. Figure 5E shows the time at which the average predictive performance was highest for each 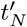. Figure S5 shows the full generalisation behaviour of the decoder.

To predict the time at which the agent would be at a particular location in Figure 5C (right), we analysed every state *s* separately, and then averaged over all *s*. For each *s*, we identified trials where the RNN passed through *s* exactly once. We trained a decoder to predict the time at which *s* was visited and computed the test performance as a function of the true time at which *s* was visited. As before, all decoders were trained in crossvalidation across current agent location.

#### Comparisons of RNN performance and efficiency

In Figure 5F-G, we compare three classes of RNNs, which were trained on either the reward landscape task, the moving goal task, or the static goal task with a goal that changed between trials. The performance of each RNN was quantified in each of the three tasks as the probability of choosing the optimal first action. We also computed the average parameter magnitudes of all the networks, 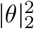. Finally, we computed the average firing rate of each network in the static goal task as 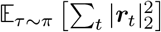. All other RNN analyses apart from Figure S7 used networks trained on the reward landscape task.

#### Subspace identification

Prior work has investigated the extent to which working memory subspaces are orthogonal (Xie et al., 2022; Dorrell et al., 2024). However, we are primarily interested in the *dynamics* between subspaces, which are easier to interpret in an orthonormal coordinate system. We therefore asked whether there exists a set of orthogonal subspaces that predict the location of the agent at every time in the future. To do so, we first simulated 6,000,000 trials and collected pairs of neural activity at time *t* and location at *t*^*′*^(*δ*) separately for each *δ*. We did this in two different ways. To estimate ‘planning’ subspaces, subspace *δ* was defined as a decoder that predicts location at time *t*^*′*^(*δ*) = *δ* from neural activity at times *t*∈ { −2, −1}. To estimate ‘execution’ subspaces, subspace *δ* was defined as a decoder that predicts location at *t*^*′*^(*δ*) = *t* + *δ* from neural activity at any *t* ≥ 0.

We then defined a predictive distribution parametrised by 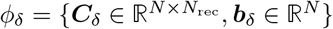 for each *δ*:

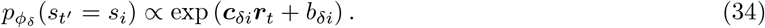

Finally, we minimised an objective function that combines the accuracy across *δ*s and the overlap between subspaces:

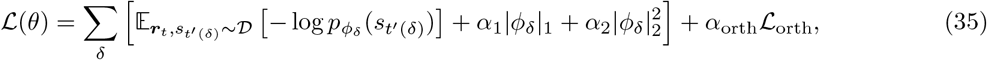

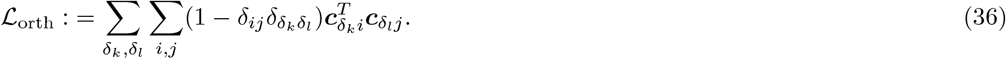

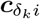 indicates the parameter vector that predicts being at location *s*_*i*_ at a delay of 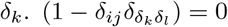 if both *i* = *j* and *δ*_*k*_ = *δ*_*l*_, and 1 otherwise. The parameters were optimised using ADAM with a learning rate of 5 *×* 10^−3^ until convergence or for a maximum of 2000 iterations. We used *α*_1_ = 10^−4^, *α*_2_ = 10^−3^, and *α*_orth_ was annealed from 0 to 2 × 10^−3^ over 500 iterations. These hyperparameters were chosen because they resulted in a good approximation to the ‘true’ subspaces in the handcrafted STA.

For the analyses in Figure S6, we considered neural activity from our 6,000,000 simulated trials separately for each time-within-trial. We then performed PCA on neural activity across all trials for each time point. We analysed both how much variance was explained by the top 80 PCs, and how much variance was explained by each of the five ‘planning’ and ‘execution’ subspaces defined by ***ĉ***_*δi*_ for *δ* = [0, 1, 2, 3, 4].

#### Estimating effective connectivity

To compute the effective connectivity between representations of different points in spacetime, we projected the learned network parameters into a coordinate system defined by the parameter vectors 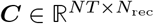. Each row of ***C*** is a normalised vector ***ĉ***_*δi*_ that predicts a particular point in spacetime. In this coordinate system, the input weights are given by 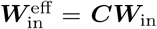; the output weights by 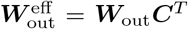; and the recurrent weights by 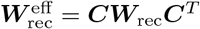. To analyse weights between ‘adjacent’ subspaces, we averaged the blocks of 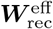 that corresponded to weights from any subspace *δ* to *δ* + 1 and from any subspace *δ* to *δ* − 1. To avoid our analyses being biased by the fact that the networks were trained with supervised learning to predict optimal locations that could only be adjacent, we did not include the weights between subspaces 0 and 1 in this average.

To quantify the similarity between the recurrent weights in this spacetime coordinate system and different order adjacency matrices for the environment, we computed point-biserial correlations. The Δ^th^ order adjacency matrix ***A***_Δ_ ∈ ℝ^*N×N*^ was defined as a binary matrix with elements equal to 1 for pairs of states that can be reached from one another in Δ actions, and 0 for all other pairs of states. The 0^th^ order adjacency matrix was defined as the identity matrix.

#### Perturbation analyses

For the perturbation analyses in Figure 6F-H, we constructed an environment where the reward function had elements (i) *R*_*ts*_ = 1.0 for (*t, s*) ∈ {(0, 0), (1, 1), (2, 2), (3, 6), (4, 10), (5, 10), (6, 10)}, (ii) *R*_*ts*_ = 0.7 for (*t, s*) ∈ {(1, 4), (2, 8), (3, 9)}, and *R*_*ts*_ = − 1.0 for all other points in spacetime. The RNN reliably converged to a representation of the optimal path. We first ran the network dynamics for 10 environment iterations without perturbation after the end of the normal planning period. We then continued to run the network dynamics for 10 environment iterations with a bias term defined by 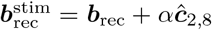, where *α* is the stimulation strength. The perturbed representation was defined as the representation at the end of this perturbation period. The two example representations in Figure 6G used *α* = 0.3 (‘weak’) and *α* = 10 (‘strong’). The quantification in Figure 6G used a range of *α* from 0 to 10 for the RNN, and from 0 to 500 for the handcrafted STA. The control analyses (grey lines in Figure 6G and Figure S9C) were performed by running the same analysis on the RNNs, but with stimulation of the same magnitude in a random direction in neural state space. For the analysis in Figure 6H, we also removed the perturbation and ran the network dynamics for 10 environment iterations after the end of the perturbation period. All analyses in the main text focused on representations in the space of implied future trajectories, 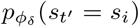. The ‘representational change’ was quantified as the L1 norm of the difference from the spacetime representation at the end of the normal planning period. See Figure S9 for an analysis of the raw firing rates.

#### RNNs trained in changing environments

For the analyses in Figure 7, we trained another set of RNNs in a version of the reward landscape task where the transition structure changed between trials. The locations of all walls in the environment were provided as a binary input to the agent. For the performance comparison in Figure 7B (top), we evaluated the performance of these networks in the environments that the ‘single structure’ networks had been trained on.

The effective connectivity in Figure 7C-D was computed as in the RNNs trained on a single structure. For these analyses, we identified the subspaces from 1,000,000 trials in a single environment, and we repeated this analysis across 30 different environments. We computed similarities between (i) the effective connectivity estimated in an environment and the adjacency matrix of the same environment, and (ii) the effective connectivity and the adjacency matrix from a different control environment. In Figure 7E, the ‘subspace similarity’ was defined as the correlation between the set of vectors that define the subspaces, averaged over all points in spacetime. We computed the similarity between (i) subspaces identified from two sets of independent trials from the same maze, and (ii) the same number of trials from two different mazes. Recall that the effective recurrent weights between subspaces are given by 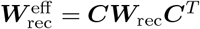. By using different subspaces in different environments, the RNN changes ***C*** between environments, which changes the effective connectivity between pairs of future subspaces (Figure 7E).

For the analyses in Figure 7G-H, we repeated the subspace identification procedure, but with two notable differences. First, we used trials across many environments to find generalised directions in neural state space that predict the future in *any* environment. Second, we defined the objective function in terms of future *transitions* 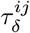 instead of locations. Figure 7G quantifies the effective connectivity between future transitions in consecutive subspaces (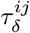 and 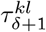) that are either ‘consistent’ (*k* = *j*), ‘adjacent’ (*j≠ k* but *s*_*i*_ and *s*_*k*_ are adjacent in an environment with no walls), or any other transitions. In Figure 7H, we first projected the input corresponding to a given wall location *w*_*ij*_ onto each subspace and normalised the projection within each subspace. We then calculated the dot product between this projection and the normalised directions in neural state space that predict either (i) transitions between *s*_*i*_ and *s*_*j*_, (ii) transitions from *s*_*i*_ or *s*_*j*_ to some other state *s*_*k*_, or (iii) transitions that do not originate at *s*_*i*_ or *s*_*j*_. Projection magnitudes were averaged over transitions within each of these three groups, then across subspaces, and finally the mean and standard deviation were computed across 5 RNNs.

## Supplementary figures

**Figure S1.**
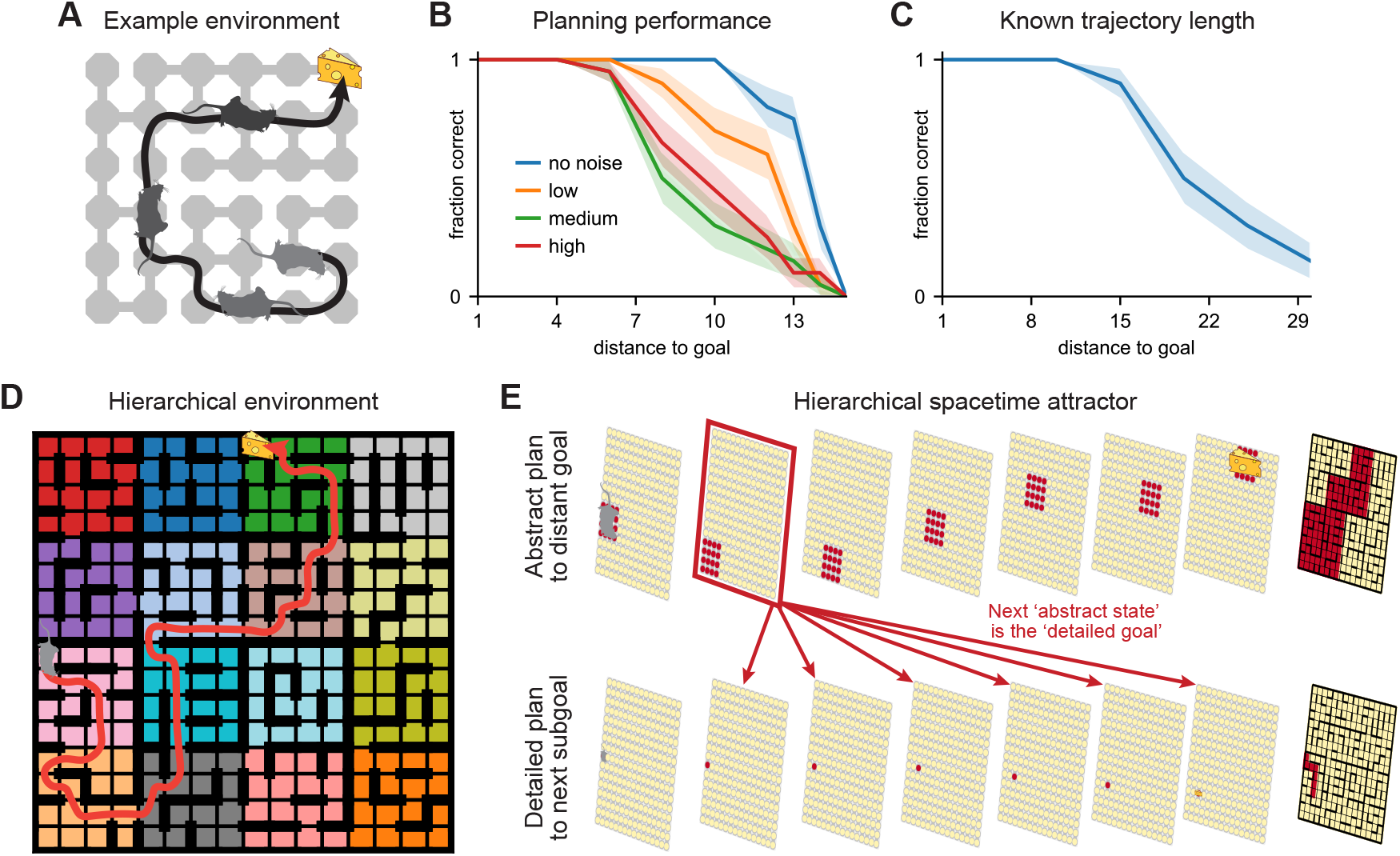
STA performance for different planning horizons. **(A)** We considered different 6×6 mazes with a single static goal location (cheese). We then quantified how reliably the STA inferred the shortest path to the goal as a function of the initial distance from the goal (mice). **(B)** Fraction of problems where the STA infers the shortest path, as a function of the distance from the goal. This analysis was repeated for four different noise levels: (i) No noise (blue). (ii) The baseline noise used in the main text (Methods; yellow). (iii) Twice the baseline noise. (iv) Three times the baseline noise. All simulations used an input strength of *β* = 6 (Methods), and representations were analysed after 1500 network iterations (30 time constants) to ensure convergence. Shading indicates standard error across 20 different environments. When the STA fails to infer the shortest path, the predominant failure mode corresponds to representations of ‘discontinuous’ paths that move immediately from the start location to the goal. This can happen if the reward inputs dominate over the recurrent signal, which decays with planning depth for intermediate locations. It can also happen if noisy connectivity leads to spurious connections between distant locations in adjacent subspaces. **(C)** The failure mode of discontinuous representations can be mitigated if the distance to the goal is known *a priori*. In this simulation, reward input was only provided beyond the first time at which the goal is reachable in a 10 *×* 10 maze with no noise. A higher input strength of *β* = 20 can then be used, and simulations were run for 300 time constants to ensure convergence. The STA reliably plans trajectories consisting of 15 actions, and it sometimes infers shortest paths that are 30 actions long. **(D)** We do not think planning depth is a major limitation of the STA as a model of human and animal behaviour, because we rarely plan more than 6-7 steps ahead at a single level of abstraction (van Opheusden et al., 2023). Instead, long-horizon planning can be solved hierarchically (Eckstein and Collins, 2020). In this example, an agent has to plan a trajectory of 40 actions (red arrow) in a large 16 *×* 16 environment. However, the environment has been grouped into 16 separate 4 *×* 4 regions (colours). **(E)** We implemented a simple hierarchical spacetime attractor. The first ‘abstract’ STA knows the transition structure between *regions*, and it infers a sequence of seven regions to the goal. The second ‘detailed’ STA receives the boundary states of the inferred next region as a reward input (red arrows). It infers a sequence of seven *primitive states* to the next region. Once the second region is reached, the abstract STA updates its representation. This allows the detailed STA to plan its way to the third region. Hierarchy enables planning of trajectories that are exponentially long in the depth of the hierarchy, and therefore in the number of neurons.

**Figure S2.**
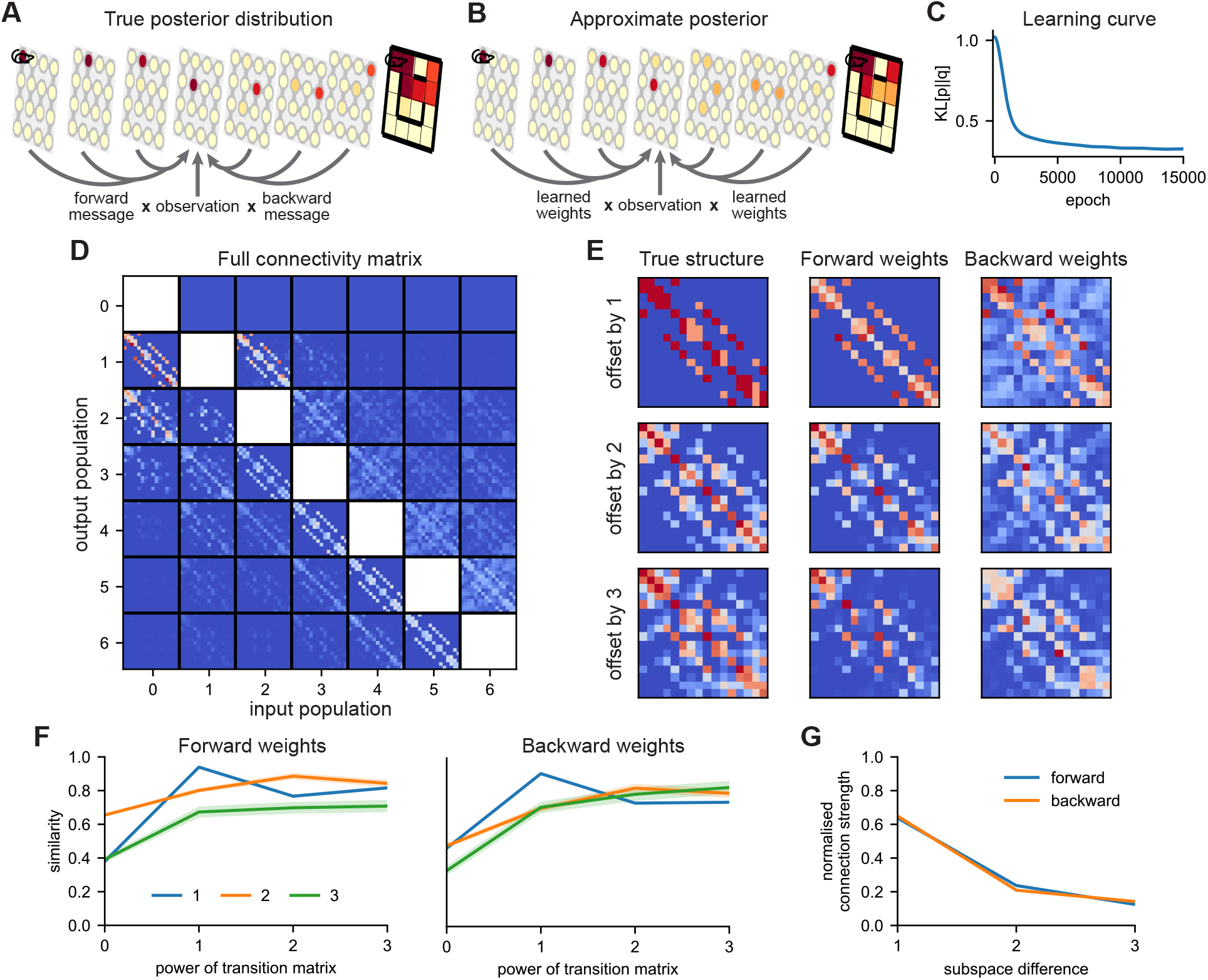
Recurrent weights optimised for planning-as-inference. **(A)** Planning can be implemented as an inference process over future locations. The posterior at each point in time is given by the product of a ‘forward message’, a ‘backward message’, and a ‘likelihood’ that reflects the reward at different points in space and time. **(B)** We trained a network with this product-of-messages architecture to learn fixed points that approximate the true posterior (Derivation). The fixed points are given by 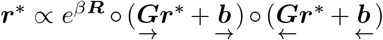, where {***G}*** are learnable parameters and *e*^*β****R***^ is the likelihood. **(C)** Average KL divergence between the true and approximate marginals over the course of learning. **(D)** Learned connectivity between all pairs of subspaces. The lower triangular blocks correspond to 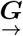, and the upper triangular blocks to 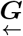. **(E)** Left: 1st, 2nd, and 3rd powers of the transition matrix. Centre: average forward connectivity between populations separated by 1, 2, or 3 actions. Right: average backward connectivity between populations separated by 1, 2, or 3 actions. **(F)** Correlation between (i) the average connectivity between subspaces separated by Δ actions (lines; legend), and (ii) different powers of the transition matrix (x-axis). **(G)** Average normalised connectivity strength as a function of the number of actions separating two populations, evaluated separately for 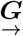 (blue) and 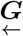 (orange). This analysis excluded connections to and from the *δ* = 0 population, which has a stronger likelihood term than all other populations. Lines and shading in (F) and (G) indicate mean and standard error across 5 networks trained in different environments.

**Figure S3.**
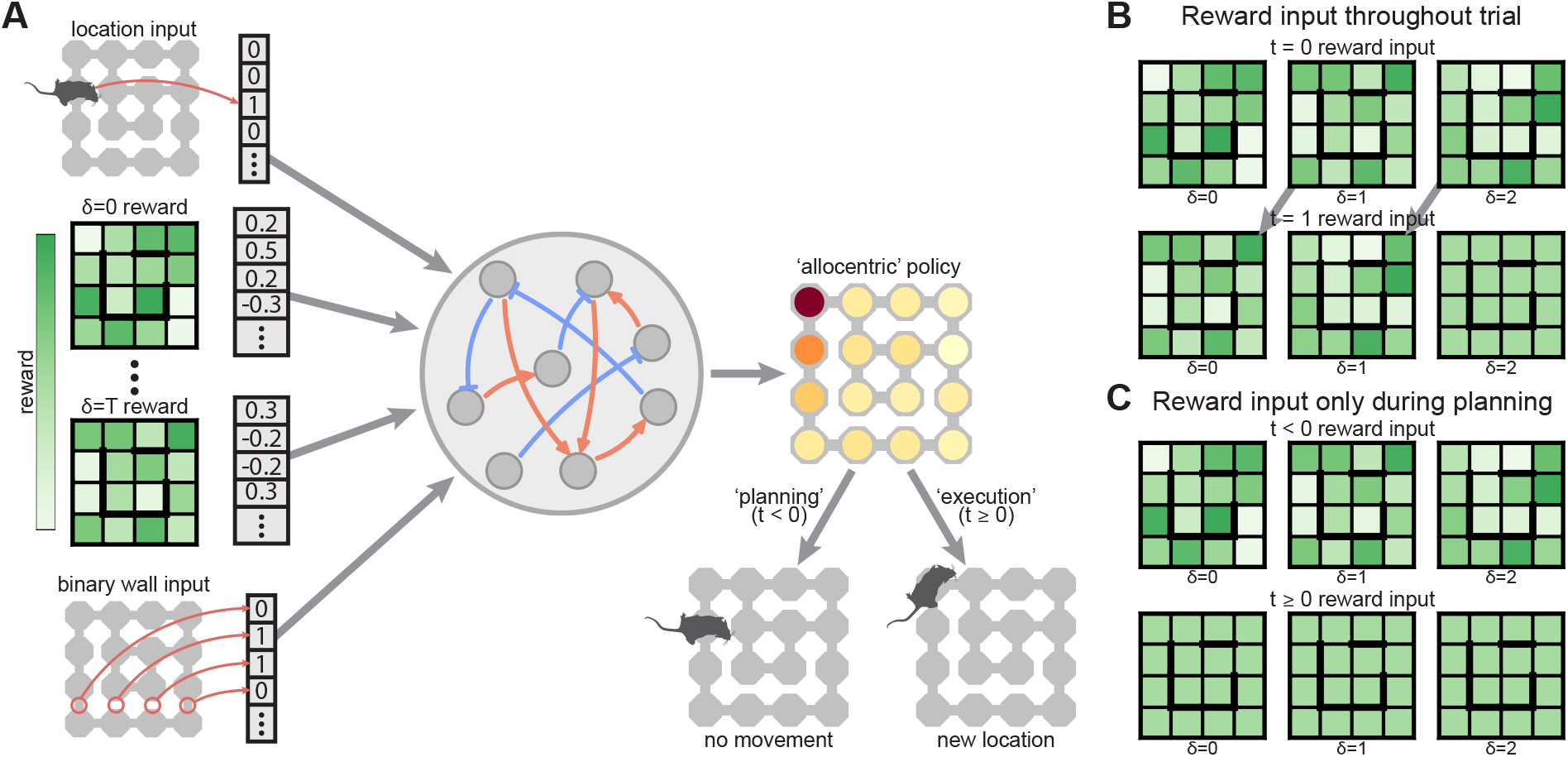
Schematic illustration of model inputs and outputs. **(A)** The RNNs and STA received three different kinds of inputs: (i) the current agent location; (ii) the scalar reward available at every location at every time in the future; and (iii) a binary vector indicating the presence or absence of barriers between otherwise adjacent states. The wall input was provided to all models, but it is only relevant for the RNNs trained in Figure 7, where the environment structure changes across trials. The output was an ‘allocentric’ policy, indicating the desirability of moving to each location in the environment. During ‘planning’ (*t <* 0; bottom left), the recurrent dynamics continued to run with no effect on the environment. During ‘execution’ (*t* ≥ 0; bottom right), the policy was normalised over locations adjacent to the current location before sampling an action. **(B)** For all analyses of the handcrafted STA and the RNN in Figure S10, reward inputs were provided continually and in ‘relative time’. In other words, a consistent input channel indicated what the reward would be at a particular location in *δ* actions. **(C)** All other RNNs were trained in a ‘working memory’ setting, where reward inputs were only provided during the initial ‘planning’ phase. During subsequent execution, all reward inputs were set to 0.

**Figure S4.**
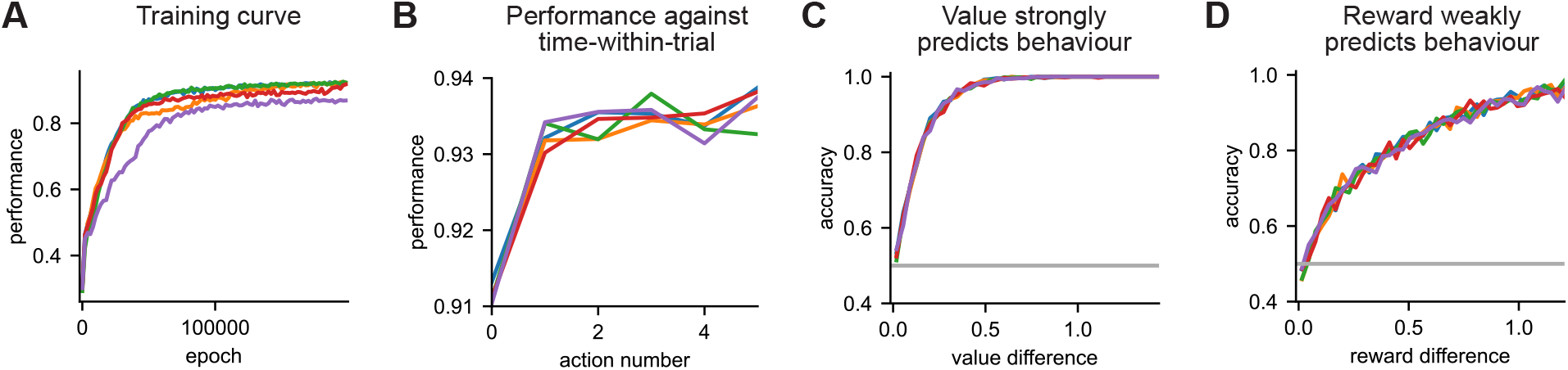
RNN performance during and after training. Each line in this figure corresponds to one of the five RNNs that were used for analyses in the main text. **(A)** Performance over the course of training, averaged over all actions within each trial. **(B)** Performance at the end of training as a function of the action number within the trial. When assessing performance at time *t*, trials were only included that had optimal choices up to time *t* − 1. **(C)** Probability of choosing the action with highest value as a function of the value difference between the two actions with highest value. This analysis shows that errors are only made then the optimal action is close in value to the second best action. **(D)** Probability of choosing the action with highest reward as a function of the reward difference between the two actions with highest reward. Reward is less predictive of behaviour than value, confirming that the RNNs compute long-term value rather than relying on greedy reward.

**Figure S5.**
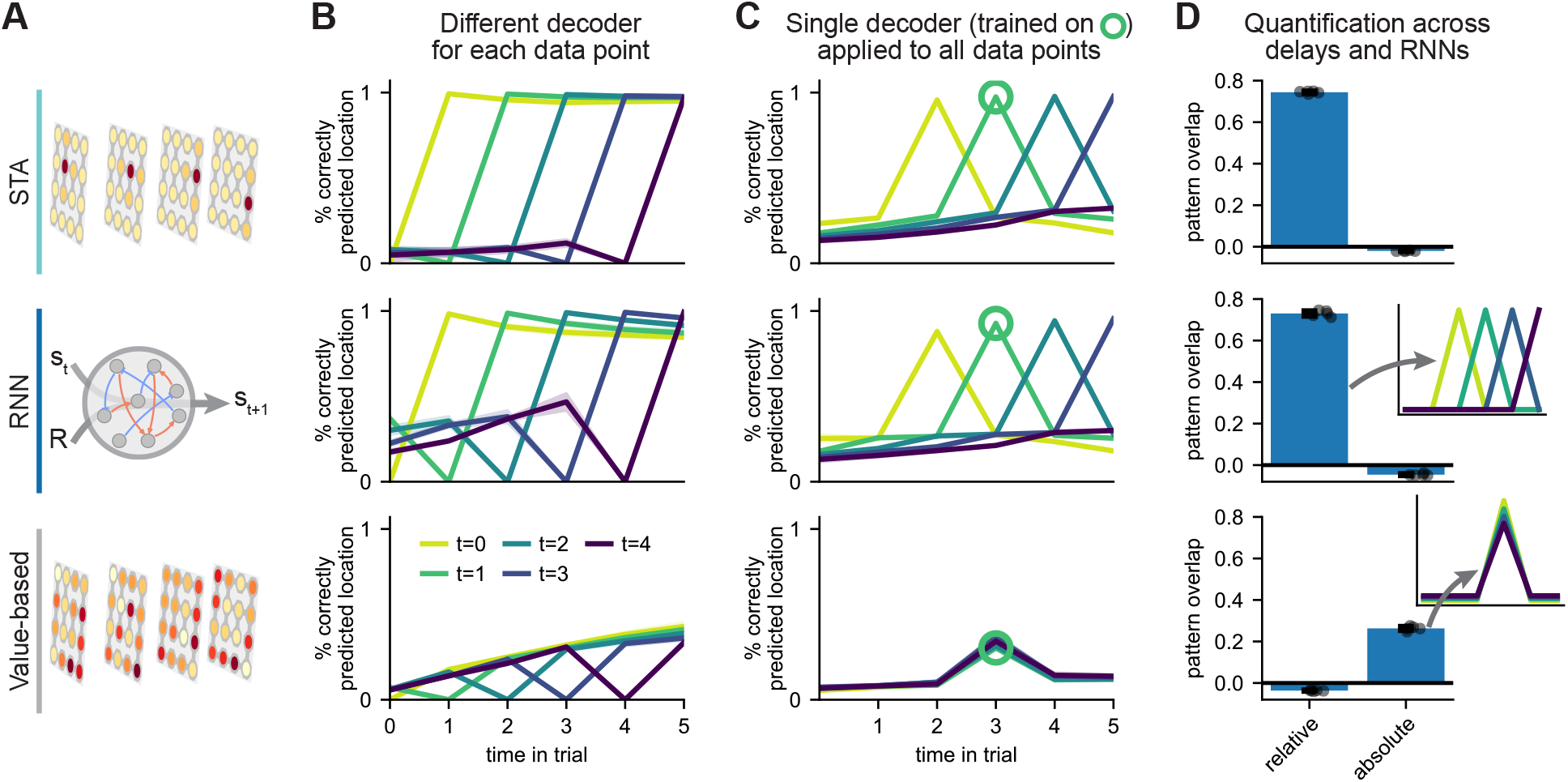
Additional analyses of learned RNN representations. **(A)** We compare a spacetime attractor; an RNN trained on the reward landscape task; and an agent that analytically computes a full spacetime value function. The value-based agent computes an optimal policy from ‘neural activity’ containing (i) the value function, (ii) the current location, and (iii) the time-within-trial (Methods). **(B)** Decoding accuracy of agent location at different times (x-axis) from neural activity at every other time (lines; legend). All decoders were trained in crossvalidation across the current agent location (Methods). This is why the accuracy is zero when decoding location from activity at the same time. **(C)** We trained a single decoder to predict location at time 3 from neural activity at time 1 (green circle). The same decoder predicts location at time *t* + 2 (x-axis) from neural activity at any other *t* (lines). **(D)** Similarity of decoding patterns to idealised representations of future location in ‘relative’ or ‘absolute’ time (schematics).

**Figure S6.**
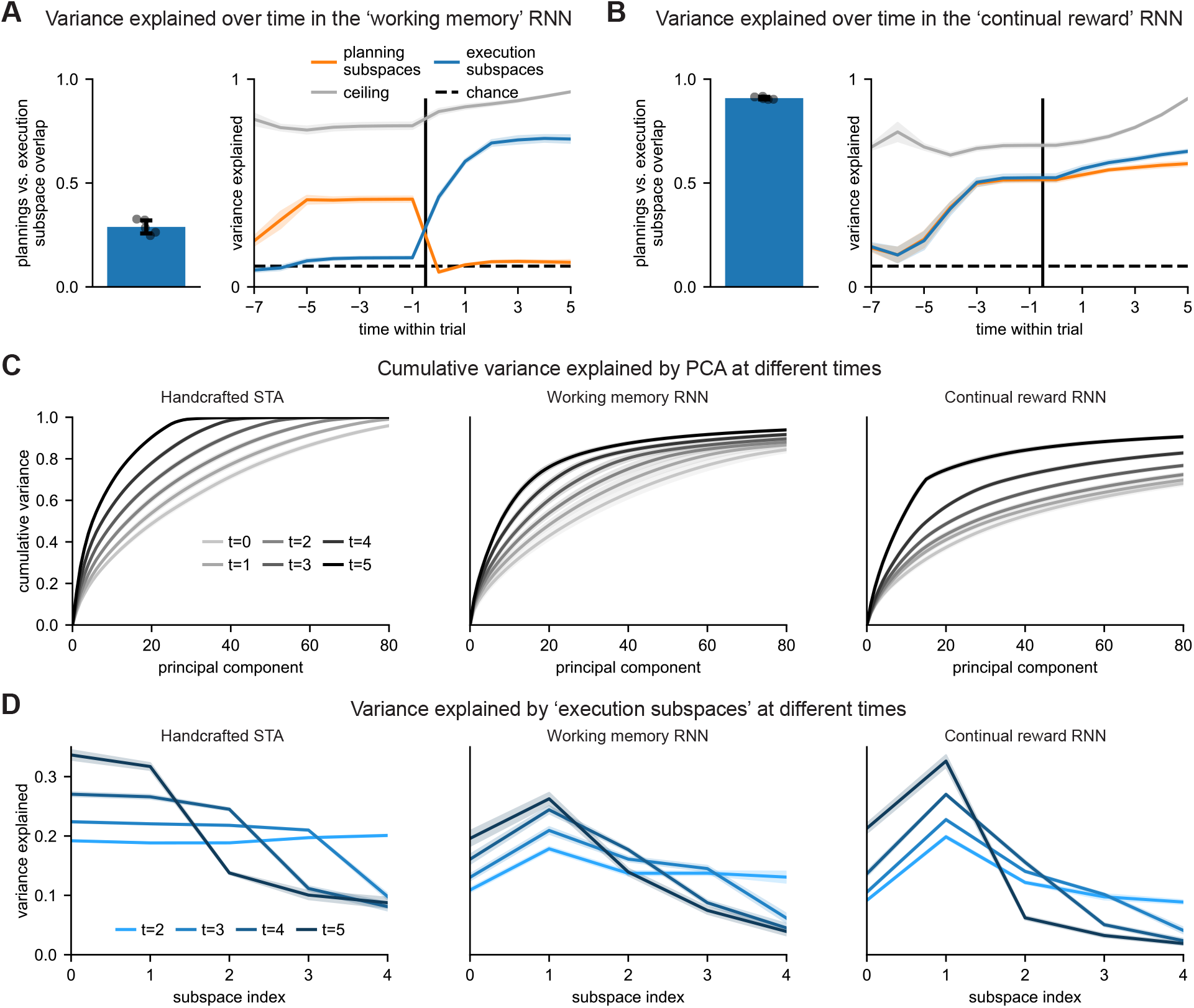
Characterisation of subspaces learned by the reward landscape RNN. We analyse the orthogonal subspaces representing future behaviour that were used to characterise RNN connectivity and dynamics in Figure 6. **(A)** Left: Empirical overlap between future-coding subspaces estimated during the planning and execution periods of the RNN. The RNN uses separate subspaces for computation of a plan and subsequent execution. Right: Variance explained by the five 16-dimensional subspaces that predict the current and next four agent locations, estimated during planning (orange) or execution (blue). The variance explained was quantified separately at each time-within-trial, which shows information transfer from the planning to the execution subspaces at movement onset (vertical line). The grey line indicates the variance explained by the top 80 PCs, which were computed separately for each time-within-trial and therefore represent a strict upper bound. **(B)** As in (A), now for a network trained with continual reward input throughout the trial instead of only during planning (Figure S3; Figure S10). **(C)** Left: cumulative normalised variance explained by the first 80 principal components of the handcrafted STA (x-axis) at different times during the execution period (lighter to darker lines). The representation becomes lower dimensional throughout execution, because the plan-to-go becomes progressively shorter. This was also true in the task-optimised RNNs (centre & right). The RNN representations were generally high-dimensional, and they had participation ratios of 50-100 during late planning and early execution. **(D)** Left: normalised variance explained by each empirical 16-dimensional execution subspace in the handcrafted STA (x-axis) at different times during the execution period (lighter to darker lines). All subspaces are active during early execution, while only the ‘immediate future’ subspaces are active during late execution. A similar trend was seen in the task-optimised RNNs (centre & right).

**Figure S7.**
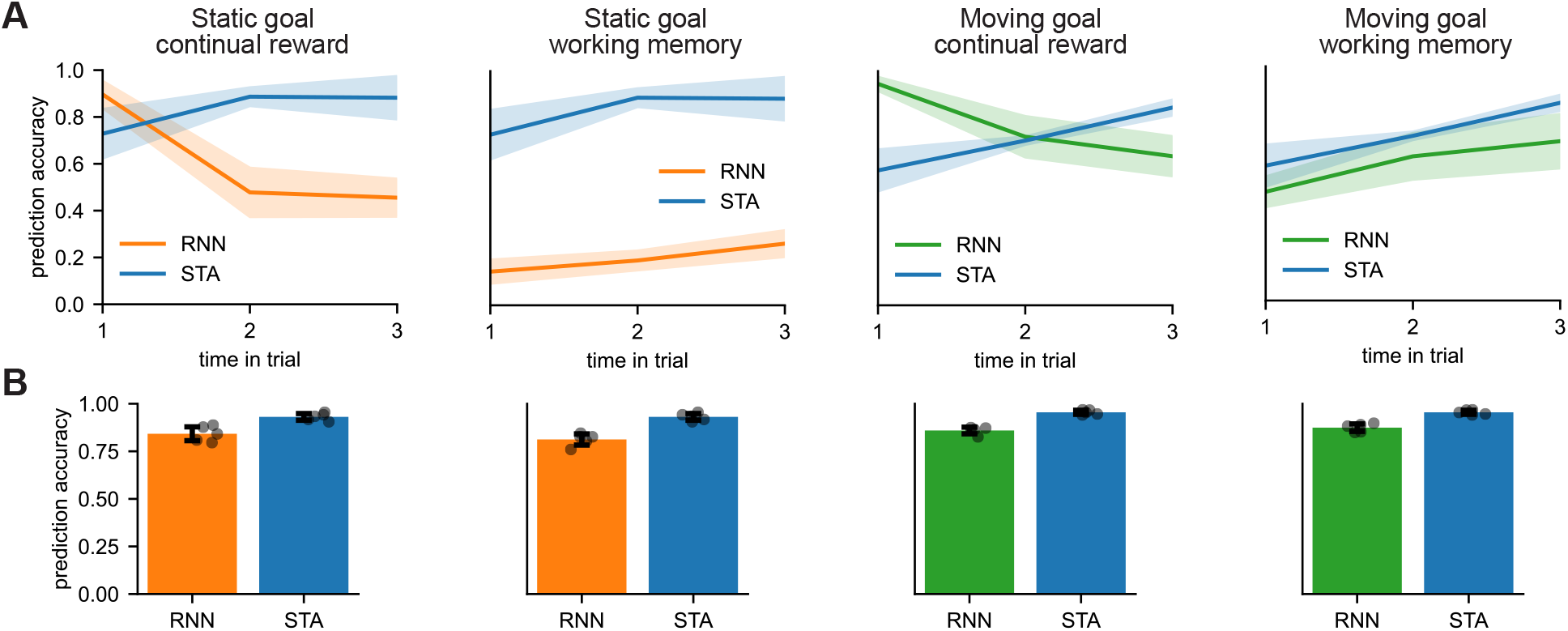
RNNs trained on simpler tasks do not learn spacetime representations. In this figure, we analyse the representations of four RNNs trained on all combinations of (i) the static goal or the moving goal task, and (ii) reward input throughout the task (‘continual’) or reward input only during the planning phase (‘working memory’). For all analyses in this figure, we only included trials where an optimal agent would intercept the goal in 3 to 6 actions. **(A)** Decoding accuracy for agent location at different times (x-axis) from neural activity at the end of the planning period. Decoders were trained in crossvalidation across the current agent location. Only the RNN trained on the moving goal task in a working memory setting learns a generalisable representation of the future. This network is also unlikely to have learned a full STA, since it fails catastrophically on the reward landscape task (Figure 5F). Note that the decoding accuracy generally *increases* slightly for the true STA as a function of time-within-trial. This is because the navigation tasks have strong correlations between consecutive positions, which leads to overfitting on the training data. This overfitting is less prominent later in a trial, where the space of possible locations conditioned on the current location is larger. **(B)** In this analysis, we trained a decoder to predict whether the agent would be at a particular location at *any* time in the trial from neural activity at the end of planning. Binary decoders were trained for each possible future location in crossvalidation across the current agent location. Bars indicate the average predictive accuracy across all binary decoders and current locations. The simpler networks learn representations of whether they will be at a given location at some point in the future.

**Figure S8.**
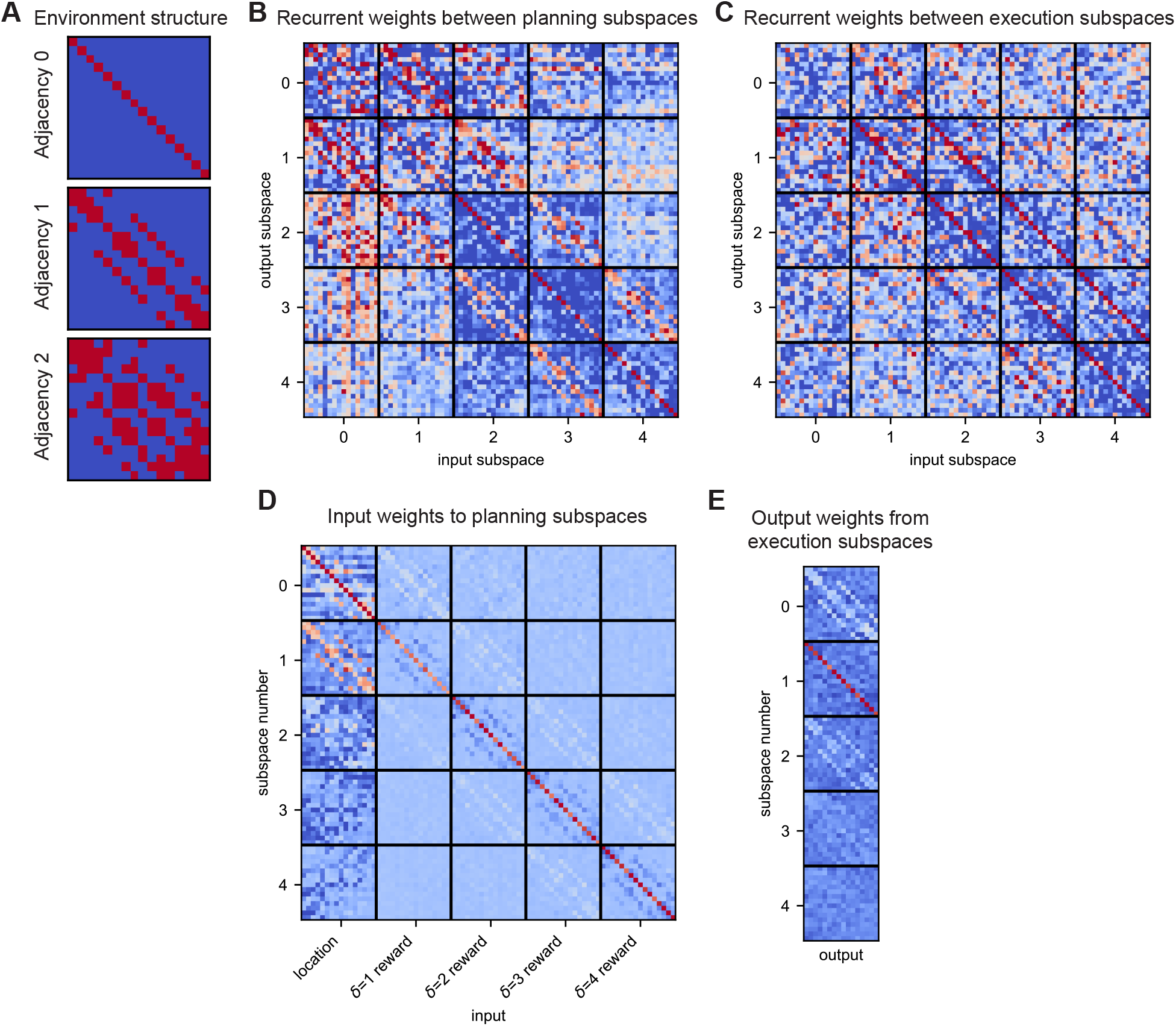
Parameters learned by the reward landscape RNN. Network weights are projected into an orthonormal coordinate system with axes that maximally predict future locations. All weight matrices are for a single example RNN, since the environment differs between networks, and the connectivity is therefore slightly different. **(A)** Structure of the environment that the RNN was trained in, illustrated as the 0^th^ order adjacency matrix (the identity matrix), the 1^st^ order adjacency matrix, and the 2^nd^ order adjacency matrix **(B)** Recurrent weight matrix estimated during the planning period, which resembles the environment adjacency matrix in the off-diagonal blocks. **(C)** Recurrent weight matrix estimated during the execution period, which has an additional ‘feedforward’ component that transfers information from later to earlier subspaces. We expect this component of the connectivity matrix to help implement the conveyor belt dynamics identified in Figure 5E (Supplementary Note). **(D)** Input weight matrix estimated during the planning period. The ‘current’ subspace receives location input, and future subspaces receive reward information corresponding to the appropriate time. **(E)** Output weight matrix estimated during the execution period. The policy is read out from the ‘immediate future’ subspace.

**Figure S9.**
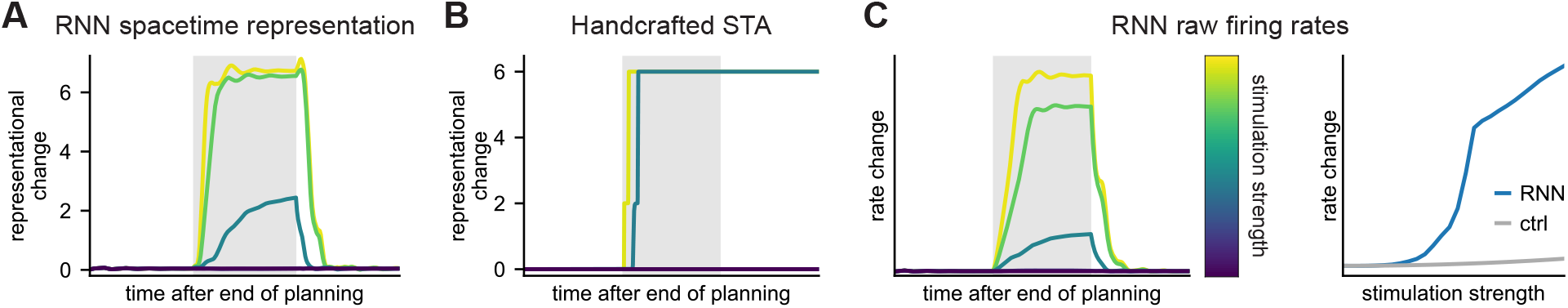
Additional analyses of attractor dynamics. **(A)** Change in implied spacetime representation over time in the trained RNN for different perturbation strengths (reproduced from Figure 6H). **(B)** Change in spacetime representation over time in the handcrafted spacetime attractor for different perturbation strengths. In contrast to the trained RNN, the ‘low value’ path is a fixed point of the perturbation-free network dynamics in the handcrafted network. At the end of a sufficiently strong perturbation, the representation can therefore stay in this new fixed point. **(C)** Change in RNN firing rates for different perturbation strengths. While the change in implied spacetime representation completely saturates with perturbation strength, the change in firing rates continues to increase with perturbation strength. This is expected because the network has a non-saturating ReLU nonlinearity. Small external perturbations are still quenched when quantifying the change in representation using the raw firing rates instead of the implied spacetime representation.

**Figure S10.**
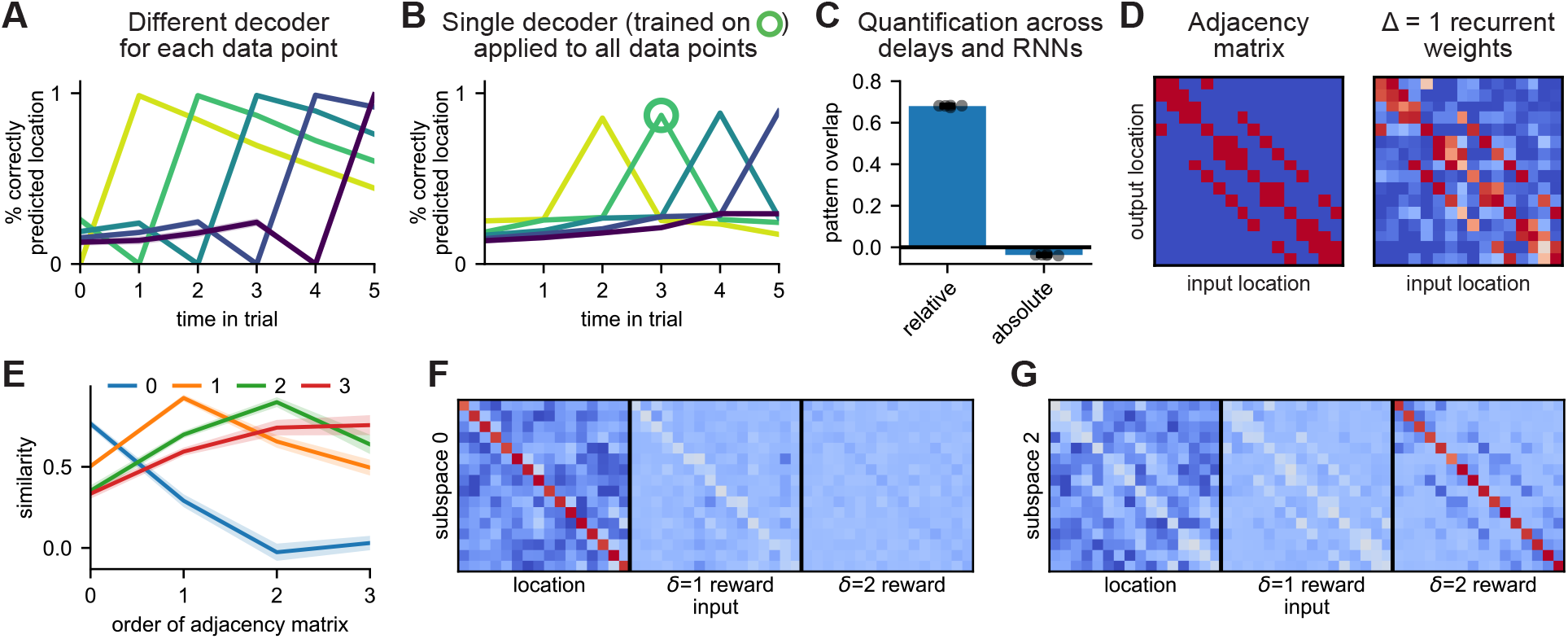
RNNs trained with continual reward input also learn spacetime attractors. In the main text, we focused on an RNN that was trained with reward input provided during an initial ‘planning phase’, while no information was given about the reward function during subsequent ‘execution’. In this figure, we perform some of the same analyses on an RNN that receives reward input throughout the entire task. In this setting, the task could in theory be solved using a ‘feedforward’ strategy that does not rely on recurrent dynamics at all. However, the RNNs still learn a spacetime attractor-like solution. **(A)** Decoding accuracy of agent location at different times (x-axis) from neural activity at every other time. Each line corresponds to predictions from neural activity at a different time in the trial from *t* = 0 (yellow) to *t* = 4 (blue). Decoders were trained in crossvalidation across the current agent location. **(B)** We trained a single decoder to predict location at time 3 from neural activity at time 1 (green circle). The same decoder predicts location at time *t* + 2 (x-axis) from neural activity at any other *t* (lines). **(C)** Similarity of decoding patterns to idealised representations of future location in ‘relative’ or ‘absolute’ time (Figure S5). **(D)** The average recurrent weights between subspaces separated by a single action resemble the adjacency matrix of the environment. **(E)** Correlation between (i) the average connectivity between subspaces separated by Δ actions (lines; legend), and (ii) different order adjacency matrices (x-axis). **(F)** Input weights to the ‘current’ subspace (*δ* = 0). **(G)** Input weights to a ‘future’ subspace (*δ* = 2).

**Figure S11.**
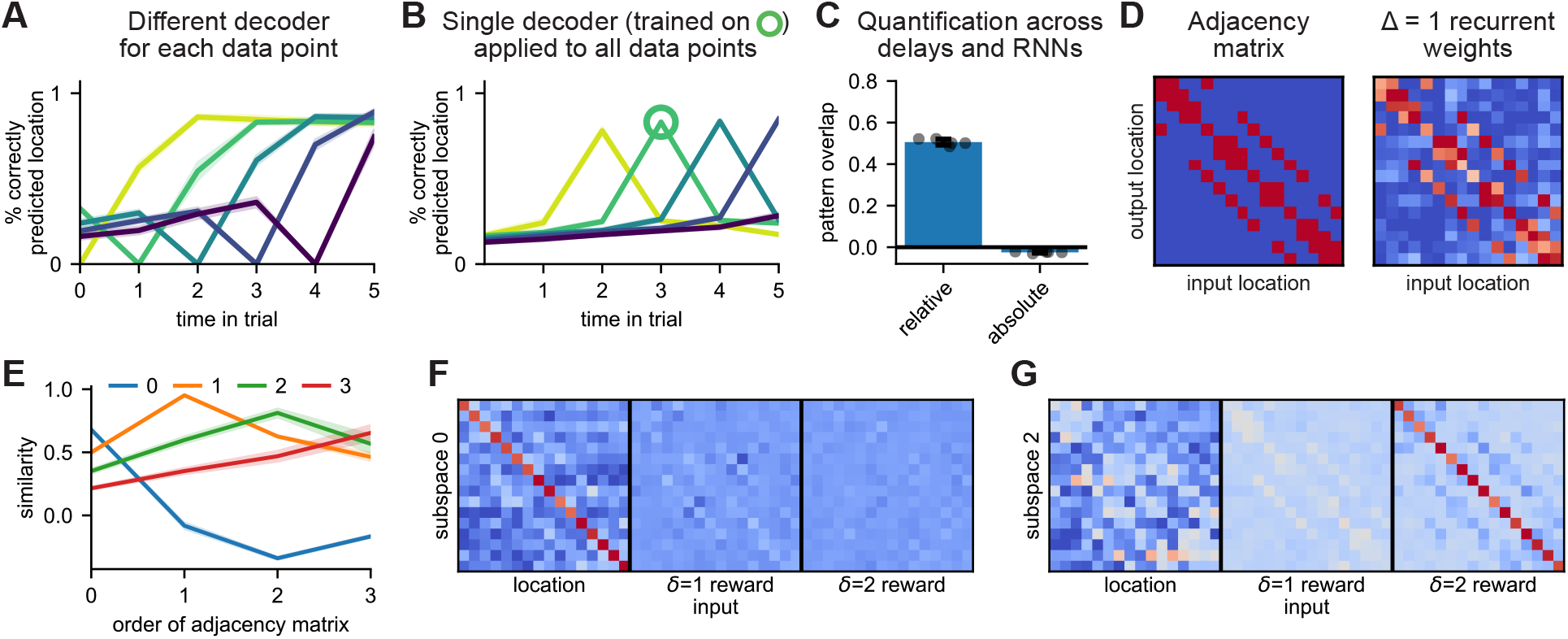
RNNs with a local action space also learn spacetime attractors. In the main text, we analysed an RNN that generated a global allocentric policy, consisting of a probability distribution over all locations that was renormalised over ‘adjacent’ locations before sampling an action. In this figure, we perform some of the same analyses on a network that outputs a ‘local’ policy in an action space consisting of ‘north’, ‘south’, ‘east’, ‘west’, and ‘stay’. This RNN also learns a spacetime attractor. **(A)** Decoding accuracy of agent location at different times (x-axis) from neural activity at every other time. Each line corresponds to predictions from neural activity at a different time in the trial from *t* = 0 (yellow) to *t* = 4 (blue). Decoders were trained in crossvalidation across the current agent location. **(B)** We trained a single decoder to predict location at time 3 from neural activity at time 1 (green circle). The same decoder predicts location at time *t* + 2 (x-axis) from neural activity at any other *t* (lines). **(C)** Similarity of decoding patterns to idealised representations of future location in ‘relative’ or ‘absolute’ time (Figure S5). **(D)** The average recurrent weights between subspaces separated by a single action resemble the adjacency matrix of the environment. **(E)** Correlation between (i) the average connectivity between subspaces separated by Δ actions (lines; legend), and (ii) different order adjacency matrices (x-axis). **(F)** Input weights to the ‘current’ subspace (*δ* = 0). **(G)** Input weights to a ‘future’ subspace (*δ* = 2).

**Figure S12.**
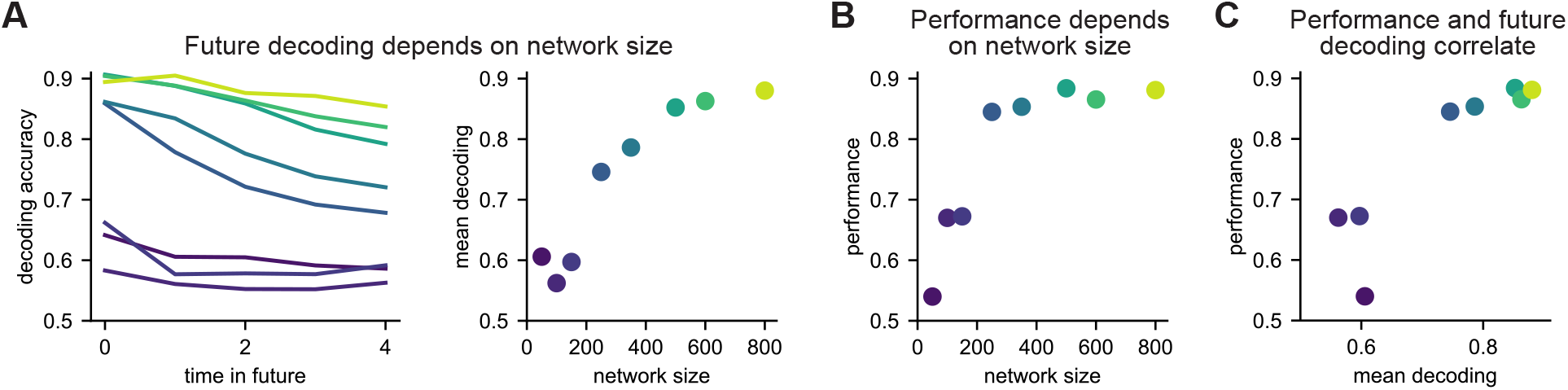
RNN representations and performance across network sizes. We trained a series of RNNs with different network sizes ranging from 50 (dark blue) to 800 (yellow) hidden units. **(A)** Networks with approximately 300 or more units learned a spacetime representation, and the future could be decoded from the hidden state of the network at the end of the planning period. **(B)** Task performance saturated as a function of network size at approximately 300 hidden units. **(C)** Task performance increased with the ability of the network to represent the entire future explicitly. These results mirror the findings of Whittington et al. (2023) that RNNs trained on working memory tasks learn a similar ‘slot-like’ solution only if the network is large enough.

**Figure S13.**
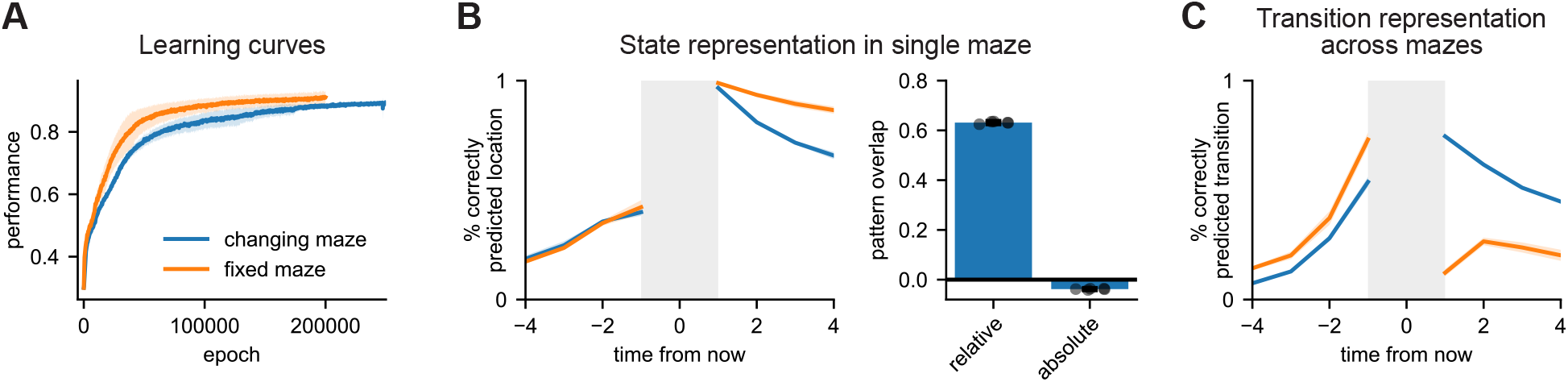
Additional analyses of RNNs trained with changing environment structure. **(A)** Learning curves of RNNs trained on the reward landscape task in a single maze (‘fixed maze’; orange) or with a different structure on every trial (‘changing maze’; blue). **(B)** We took the RNNs trained across many mazes and evaluated them in a single maze (blue). Future states could be decoded almost as well as in the RNNs trained in a single maze (orange). Additionally, the representations in each maze were more similar to idealised representations of future location in ‘relative’ than ‘absolute’ time (right). **(C)** When considering data across many mazes, future *transitions* could be decoded in a way that generalised across current transition and maze structure (blue). Such a representation did not exist in RNNs trained in a single maze (orange).

## Supplementary discussion

In this section, we discuss some of the many architectural and modelling choices that went into our work. As is the case for much work in modern computational neuroscience, the space of models was larger than we could feasibly explore fully in a single paper. In what follows, we aim to further motivate the choices that were made in the main paper and to provide additional intuition for the importance and effect of various architectural choices and hyperparameters in our work. We hope this will be useful for readers looking to better understand our work or draw inspiration from it in their own research.

### Source of reward information

In this work, we have studied the challenge of planning – computing an optimal trajectory given a particular initial state and reward function. We have therefore assumed access to ground truth rewards across all simulations. In practice, these rewards would themselves have to be computed and represented somewhere in order to provide them as an input to a spacetime attractor. We think there is no single answer to where such reward information comes from in biological circuits. Sometimes, it could come through structural knowledge – we know what ‘checkmate’ looks like on a chess board, which allows us to assign positive reward to winning positions. Other times, it could come through explicit instruction – if a reviewer asks for a new analysis, I should write the code to do it. Many times, an effective reward is inferred from complex interactions with the environment. I may have noticed that I only had one egg left after breakfast, and later remembered that it is my friend’s birthday tomorrow. Now I have to go to the grocery store before it closes at 6 pm so I can bake a cake in the evening.

In terms of neural representations, there is some evidence for goal-related coding in the hippocampus (Nyberg et al., 2022) and orbitofrontal cortex Basu et al. (2021). It is therefore plausible that the hippocampus and orbitofrontal cortex are involved goal *selection* – or latent state inference (Mishchanchuk et al., 2024) – and a spacetime attractor in medial prefrontal cortex takes a particular goal as an *input* to infer a trajectory to the goal. However, we think these putative neural representations of reward constitute only a small subset of reward inference and goal coding during natural behaviour. Reward inputs could for example also be inferred by a spacetime attractor itself. For the hierarchical planner in Figure S1E, the reward input to one STA is itself an inferred state from an STA operating at a higher level of abstraction. Alternatively, multiple STAs representing different individuals could interact during social computations. In this case, the reward input to each STA is an output of the others. Unfortunately, it is quite challenging to *study* the source of reward information in natural settings, because experimenters usually impose an explicit reward function to motivate experimental subjects. Whatever the source is, we hope it will be useful to understand how distant rewards are used to shape our immediate actions.

### Reward input format

When training RNNs with a reward function that changes in time, there are multiple different ways this could be provided as an input to the agent. The three most natural choices are:

1. A relative encoding, where a given input channel always reflects the reward in *δ* actions.
2. An absolute encoding, where a given input channel always reflects reward at time *t*.
3. A ‘working memory’ setting, where reward input is only provided during a planning period prior to the first action. At this time, the relative and absolute representations are identical, and no choice has to be made between the two.

One might expect a relative reward input to bias RNNs towards relative future representations, and ‘conveyor belt’ dynamics, and an absolute reward input to bias RNNs towards an absolute code. While RNNs trained with relative reward input did show strong relative coding of the future (Figure S10), RNNs trained with an absolute input learned somewhat mixed representations that were harder to interpret.

To avoid having to choose between these two, most of the analyses in the paper were done on an RNN trained in the ‘working memory’ setting. We also saw slightly stronger ‘spacetime’ representations in the working memory setting. In fact, Figure S10 shows that the effective planning horizon of the ‘relative’ model is only 4-5 steps, and predictive performance of future states drops off beyond that. This is presumably because a shorter planning horizon is sufficient for near-optimal performance in the task, and the regularisation encourages networks to not encode more information than necessary. The RNN therefore effectively re-plans at every step using a spacetime attractor with a depth of approximately 4.

The handcrafted spacetime attractor always received a ‘relative’ reward input. Otherwise, it would need to include either (i) an additional ‘memory’ component, (ii) a transformation from absolute to relative inputs, or (iii) a mechanism for changing the readout between subspaces. We expect that if biological agents use spacetime attractor-like dynamics, there may be settings where the code is relative and settings where it is absolute. This might depend on whether the plan is shaped around constraints in relative time (a meeting in 10 minutes) or absolute time (a meeting at 1 pm).

### Planning period

When training the RNNs, we included a planning period prior to the first action. During this period, the output of the network had no effect on the environment. We did this (i) to separate the ‘planning’ period from the ‘execution’ period in the working memory setting, and (ii) because it slightly improved performance. In the handcrafted STA, there is no distinction between planning and execution, since the inputs are the same in both cases. However, convergence can be slower before the initial action because the network starts further from a fixed point. This means that the STA can be run for fewer iterations after the first action, which effectively corresponds to a longer ‘planning phase’ followed by faster execution.

### Network iterations per action

For all RNNs, we used a variable number of planning iterations. This is because we were interested in stable representations of future behaviour, rather than networks learning to time their dynamics to initiation. In the RNNs with a continual reward input, we varied the number of network iterations between each action for the same reason. However, this is not possible in the working memory RNN, since it would not know when it had taken an action. There are two possible solutions to this problem. (i) we can provide a specific ‘action’ input that tells the network when the environment has updated. (ii) we can use a fixed number of network iterations between every action and allow the network to learn the speed of the environment. We opted for the second option to keep the inputs as simple as possible.

In the ‘continual reward’ RNNs, we suspect that the neural dynamics will approach a fixed point before each action, because a good policy must be maintained throughout the possible action times. The working memory RNNs could instead use different subspaces at different ‘phases’ of the execution dynamics, similar to findings by Stroud et al. (2023) in simpler working memory tasks. However, we did not test this hypothesis explicitly. Instead, we focused on the planning period when analysing the connectivity of the working memory RNNs to avoid potential complications due to such phase coding. When analysing the representations during ‘execution’ in Figure 5D, we used only neural activity right before each action.

### Execution-time dynamics

The fixed points of the STA dynamics are input-dependent (Figure 3), and the inputs to the handcrafted model change when it takes an action, because the location and/or future reward will be different. For this reason, the representation can automatically update to reflect the ‘path-to-go’ after each action. However, this involves recomputing the future to some extent, which is ‘wasteful’ since it has already been computed once. Additionally, the representation can get stuck in local minima because the barrier to switching between representations is non-zero. We therefore included an additional component in the STA weight matrix, which was 1 for all off-diagonal elements that implement a transfer of information to earlier subspaces. This transfer component was only active for 100 network iterations (two time constants) after each action to update the representation before settling into a new fixed point. This is similar to how the fruit fly head direction circuit uses populations of ‘shift’ (PEN-1) neurons to update the internal heading, and these PEN-1 neurons are gated by angular velocity input (Turner-Evans et al., 2017, 2020). Including an explicit shift component in the STA is not necessary for any of the main results in the paper, but it helps stabilise the dynamics.

### Noise sensitivity

The handcrafted STA is more or less susceptible to different types of noise. The model is very robust to noise added to the neural potential (‘*z*’; Methods), which can be interpreted as a spacetime distribution in log probability space. This is because non-desirable locations in spacetime can have very negative values, which are not very noise sensitive. The STA is more sensitive to noise added to the firing rates (‘*r*’; Methods), which can be interpreted as a spacetime distribution in actual probability space. This is because addition of positive noise to some location *s*_*i*_ in subspace *δ* will propagate through the adjacency matrix to all neighboring locations at time *δ* + 1. This can lead to representations that ‘teleport’ between distant locations if *s*_*i*_ is near the goal and therefore receives strong excitation either (i) directly from the reward input, or (ii) indirectly through the recurrent input from future subspaces.

The STA is sensitive to structural noise for a similar reason. Adding a small positive value to weights corresponding to elements of the adjacency matrix that are meant to be zero leads to dynamics that include a finite probability of teleporting between these distant locations. In this work, we mitigate the structural noise sensitivity by using a ‘pessimistic’ estimate of the adjacency matrix as the base weights before adding noise (Methods). In other words, we subtract a small constant from all weights to ensure that distant locations are connected with weights that are zero or negative, even though they are noisy. This may be less of a problem in biological networks, since Dale’s law ensures that synapses are either excitatory, inhibitory, or absent.

The sensitivity to both rate noise and structural noise is higher when the input strength is larger (*β*; Methods), which biases the representation more strongly towards rewarded locations. Conversely, the converged representation is more diffuse if the input strength is weaker, because there is a smaller bias towards rewarded locations. This is particularly true when planning towards distant rewards. The strength of the reward input therefore has to balance the planning depth with the susceptibility to teleporting representations. If there is no noise in the system, the input strength can safely be very large, which leads to robust performance for long planning horizons. If there is more noise in the system, the input strength should be smaller, which increases noise robustness at the expense of a shorter effective planning horizon. When training RNNs across tasks, they will naturally learn to balance the robustness of the representation with the required planning depth. Indeed, the analyses in Figure S9 suggest that RNNs learn to do so better than our handcrafted models.

### Choice of RNN learning algorithm

We used supervised learning to train all RNNs in this paper. In other words, the RNNs were trained to predict the behaviour of an optimal agent. An alternative would have been to train the networks end-to-end using reinforcement learning. We opted for the supervised setting for two reasons. Firstly, supervised learning is more stable, which leads to more robust results that are less sensitive to hyperparameters and use less compute. These features make it much simpler for others to build on our work. Secondly, we do not believe that cortical representations are learned from scratch via reinforcement learning. Instead, we think they are learned via predictive or ‘semi-supervised’ learning. The basal ganglia can then use reinforcement learning to map these representations onto actions (Blanco-Pozo et al., 2024; Zintgraf et al., 2019). We expect that training the RNNs with reinforcement learning would yield similar results, but we have not tested this explicitly.

### Convergence of RNN training

Convergence was in general fairly good, but it did vary with hyperparameters. For some combinations of regularisation strengths, the networks sometimes got trapped in local minima that corresponded to incomplete structural learning. These networks would partially learn the structure of the environment, but they would fail to learn connections between some pairs of states that were actually connected, which impaired performance. Similarly, some RNNs trained on the changing maze task erroneously ‘hard coded’ some transitions instead of having them flexibly modulated by the inputs. In general, the accuracy of the learned structure was correlated with model performance for both the networks trained in a single maze and networks trained in changing mazes. Additionally, the overall loss was higher for the networks that failed to learn the full task structure, suggesting that it is an issue of convergence rather than regularisation making the partially learned solution optimal.

### Long distance parameters in the RNN

In Figure 6 and Figure S8, we saw that the trained RNNs learn some long-range connections between subspaces separated by more than one action. This differs from the handcrafted STA, which only has connections between adjacent subspaces. However, the presence of long-range connections in the RNN is not too surprising. In particular, we expect this additional structure to stabilise fixed points corresponding to possible trajectories, since it inhibits any trajectory that includes impossible n-step transitions. It also mirrors our finding in the networks directly optimised for planning-as-inference (Figure S2). We suspect that stabilising the dynamics through weaker connectivity between all subspaces is cheaper than strong connectivity exclusively between adjacent subspaces. This may be a consequence of the L2 regularisation used to train the networks, which favours many small parameters over few large parameters. In future work, it could be interesting to explore whether the strength of long range connectivity is lower when using L1 regularisation instead. The result of such an analysis may also depend on how disentangled the spacetime representations are, which we did not explore in the present paper.

### Discrete space and time

We have discretised space and time throughout this work. This makes the models and analyses simpler, because the action space is discrete and one action always leads to a step change in the environment. However, we expect that the basic ideas extend to continuous space and time as well. In this case, neurons would still represent particular points in spacetime. Pairs of neurons would be connected as a function of their difference in preferred location in a way that reflects which locations can be reached in a unit time. This is similar to the mechanism used for angular velocity integration in ring attractors and path integration in grid attractors. If the speed of the agent can vary, it might be necessary to represent different speeds in different connections or neurons, and the desired speed at every point in time could be inferred together with the trajectory.

### Stochastic environments

We have worked with deterministic environments throughout this paper. This choice greatly simplifies the spacetime attractor, since the deterministic adjacency matrix can be built into the network connectivity. When working in a stochastic environment, we would intuitively want the connections to represent max_*a*_ *p*(*s*_*t*+1_|*s*_*t*_, *a*), which is a generalisation of the adjacency matrix for deterministic environments. We have not explored this explicitly but consider it an important extension for future work. One potential challenge in the stochastic case is that the spacetime attractor as formulated here approximates planning-as-inference under the assumption that the posterior distribution over locations factorises across time. This assumption may have more severe consequences in stochastic environments, where correlations are more important.

### Relationship to sequence working memory

The spacetime attractor is strongly inspired by representations identified for sequence working memory (El-Gaby et al., 2023; Xie et al., 2022; Chen et al., 2024; Tian et al., 2024; Whittington et al., 2023). These sequence memory tasks have notable similarities to the reward landscape task studied in this work. In particular, sequence working memory often involves presenting an animal with a sequence of 2-4 options (e.g. ‘a’, ‘b’, and ‘c’) that are sampled from a set of N possible states or actions (Xie et al., 2022; El-Gaby et al., 2023). The animal is then rewarded for repeating the sequence at response time. The task therefore has a reward function that is categorical (only one element is rewarded at a given moment in time) and changes in time (first ‘a’ is rewarded, then ‘b’…). This is very similar to the changing reward function in the reward landscape task. A notable difference is that the sequence memory tasks have no constraints on the possible transitions – any element can be produced before or after any other element. For this reason, each sequence element can be treated independently, and there is no need to pass reward or value information between subspaces. The independent sequence working memory task can therefore be seen as a special case of the reward landscape task, where (i) only one state is rewarded at each point in time, and (ii) any state can be reached from any other state, so the adjacency matrix is uniform. Interestingly, theoretical work shows that explicit representations of the future preferentially emerge for sequence working memory when correlations (or more precisely, ‘range dependence’) between subsequent elements are weak, and the space of possible sequences is therefore large (Dorrell et al., 2024, 2026). This is reminiscent of our finding that spacetime representations for planning preferentially emerge in RNNs trained on the reward landscape task, where the space of possible optimal trajectories is larger than in the static and moving goal tasks.

### Comparisons with TD and SR agents

In Figure 4, we compare the representations and performance of the STA to temporal difference learners and successor representation agents. We show that the STA solves ‘dynamic’ problems that these algorithms struggle with. This is in some sense a property of the state space as well as the decision making algorithm. It would be possible to construct TD and SR agents in a ‘space-by-time’ state space, which would be better suited to these dynamic problems. Our claim is not that this is not possible. Instead we are highlighting that most neural implementations of decision making involve representations that are ‘flat’ across time, and we use them as a comparison to show why spacetime representations can be useful. If a TD learner was implemented with a space-by-time state space, it would be able to solve tasks where the reward changes within a trial but the same pattern is seen across all trials. The SR agent with a space-by-time state space could solve problems with reward functions that change within a trial. We expect such a spacetime-SR to work well for problems with a single moving goal, since the expected occupancy under a diffusive policy correlates strongly with the shortest path. It is less clear how well it would work for the reward landscape task, where it may be be optimal to go to high-reward locations in close proximity to locations with negative reward. This would lead to low expected reward under a diffusive base policy, but high reward under the optimal policy.

In the simplest implementation, a spacetime-SR would require inversion of a matrix T ∈ ℝ^*NT×NT*^ with a memory cost of *O*(*N* ^2^*T* ^2^) and a computational cost of *O*(*N* ^3^*T* ^3^). It is possible that the structure of T could be exploited to invert it more efficiently, which would be an interesting avenue for future research. Conversely, the STA has a memory cost of *O*(*NT*) – the number of neurons in the network. The computational cost is *O*(*N* ^2^*T* ^2^) because each iteration of the network dynamics requires *T* matrix-vector products of size *N × N*, and the number of steps to convergence is roughly linear in *T*.

### Related work on sequence generation in attractor networks

Previous work has also investigated how sequences can be generated by neural networks with attractor dynamics. Classical Hopfield networks (Hopfield, 1982) have been extended by Kleinfeld (1986) and Sompolinsky and Kanter (1986) to incorporate asymmetric connectivity that drives sequence generation. In both cases, this was achieved through *delayed inputs* that allowed the system to converge to a stable representation before moving on to the next ‘memory’ in the sequence. More recent work has also derived mechanistic models of sequence generation through asymmetric connectivity (Parmelee et al., 2022) or spike-frequency adaptation (Widloski et al., 2025). Common to these models is the idea that a single population of neurons encodes a sequence through the temporal evolution of its activity, with the dynamics of the system giving rise to limit cycles. These sequential representations are distinct from the STA, where an entire sequence is encoded instantaneously as a stable fixed point of the network dynamics. This instantaneous representation of entire sequences is what allows planning to proceed through parallel message passing rather than sequential search.

### Summary

As is evident from this discussion, many choices went into this work that could have been different. We do not claim to have explored the full space of models and tasks, and we are not trying to argue that the spacetime attractor is a silver bullet for planning and decision making. Instead, we suggest that STA-like models are interesting solutions to a range of problems that are relevant to prefrontal cortex and not widely studied in systems neuroscience. However, many open questions remain, some of which we have briefly motivated here. We hope the insights from this paper will be helpful for further exploration of these questions, both experimentally and computationally.

## Derivation for approximate inference

In this section, we derive the equations used to optimise the model in Figure S2. Our goal is to implement planning-as-inference through the recurrent dynamics of a neural network. We define ***ω*** ∈ ℝ^*NT*^ as the vector of emission probabilities, *ω*_*t,i*_ = *p*(*o*_*t*_ = 1|*s*_*t*_ = *i*); ***p*** ∈ ℝ^*NT*^ as the vector of marginal posterior probabilities, *p*_*t,i*_ = *p*(*s*_*t*_ = *i*|*o*_1:*T*_); and ***T*** ∈ ℝ^*N×N*^ as the prior transition matrix, *T*_*ij*_ = *p*(*s*_*t*_ = *i*| *s*_*t*_ *™*_1_ = *j*). We use ***v***_*t*_ ∈ ℝ^*N*^ to refer to the block of vector ***v*** ℝ^*NT*^ that relates to time *t*. Exact planning-as-inference can then be implemented using the ‘forward-backward’ algorithm:

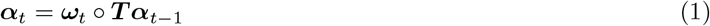

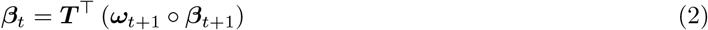

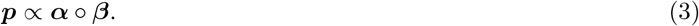

We can instead construct the matrix T ∈ ℝ^*NT×NT*^, which contains ***T*** in its lower off-diagonal blocks; the ‘boundary vectors’ 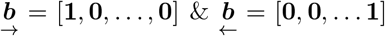; and the diagonal matrix Ω = diag(***ω***) ∈ ℝ^*NT×NT*^. This allows us to rewrite the posterior in block form:

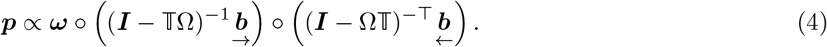

This is the product of (i) a likelihood, (ii) a forward message, and (iii) a backward message. However, the forward and backward messages include terms in Ω. This means that the effective ‘weights’ of the network have to change from problem to problem.

We take inspiration from the form of the exact posterior, but we instead rewrite the forward and backward messages as a product of (i) a set of fixed weights that generalise across problems, and (ii) the approximate marginals *q*_*t,i*_ ≈ *p*(*s*_*t*_ = *i* |*o*_1:*T*_). It is not obvious *a priori* what ‘messages’ should be sent between the different ***q***_*t*_ to make this approximation as good as possible. However, the fixed points should have the form

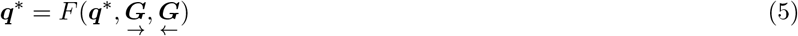

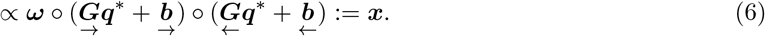

***q*** ∈ ℝ^*NT*^ is normalised separately within each time block,

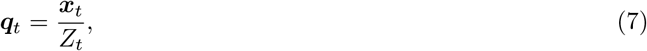

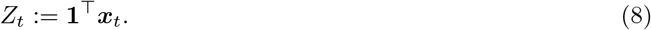

For notational convenience, we define the shorthands

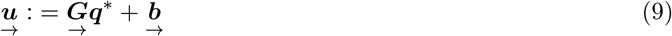

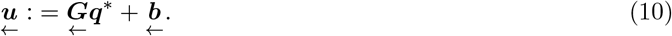

Our goal is to maximise the negative KL divergence between the true and approximate marginals,

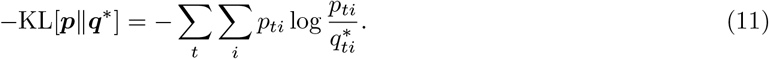

Dropping terms independent of ***q***^∗^, the objective is

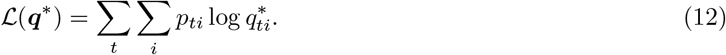

The gradient w.r.t. 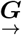 is given by the chain rule:

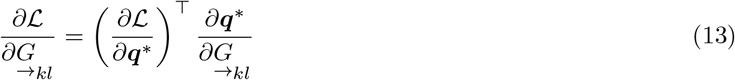

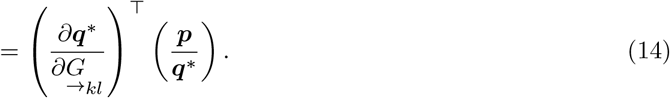

We now derive the implicit gradient of the fixed point with respect to 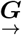. Differentiating both sides of Equation 5 gives

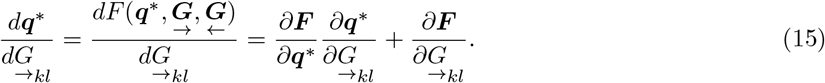

Rearranging,

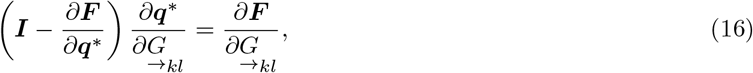

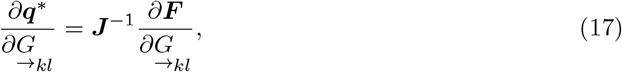

where 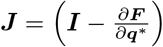. To compute ***J***, we note that:

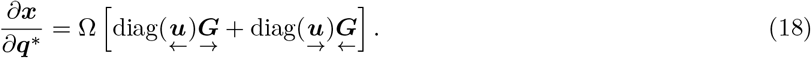

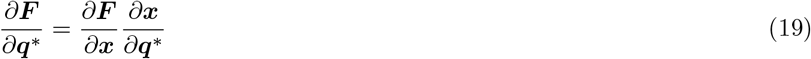

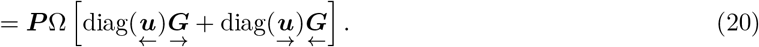

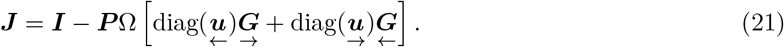

Here, ***P*** is the Jacobian of the normalisation:

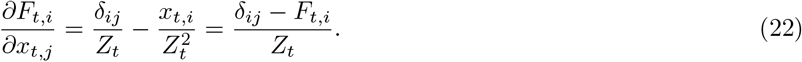

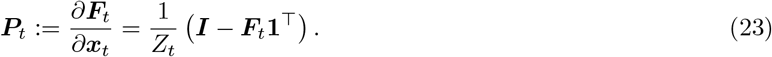

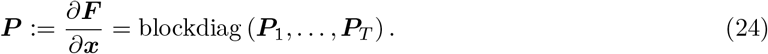

Since ***F*** depends on 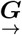 only through ***x***,

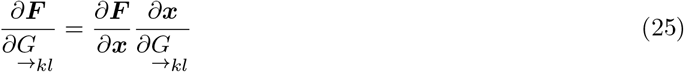

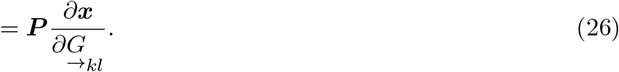

Defining ***e***_*k*_ as the *k*th Cartesian basis vector, we get

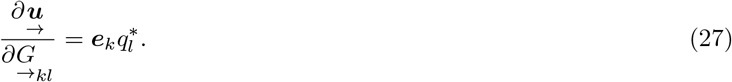

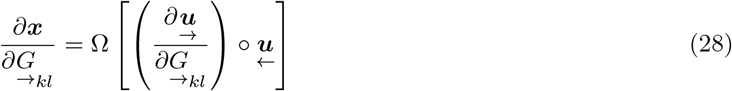

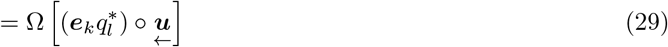

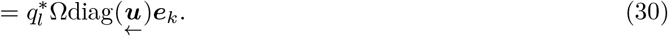

Consequently,

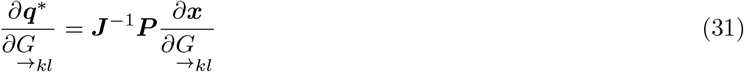

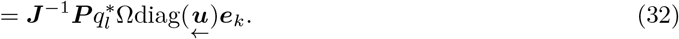

The scalar gradient of the total loss is

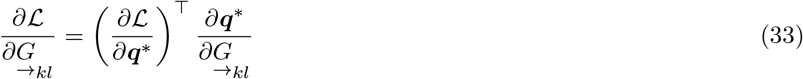

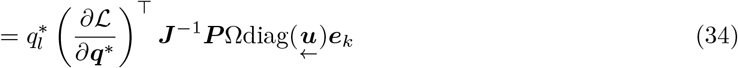

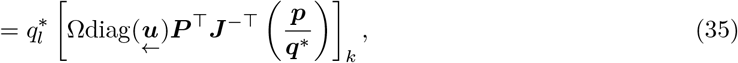

where the last line follows from taking the transpose and substituting 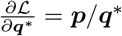. Reassembling into matrix form:

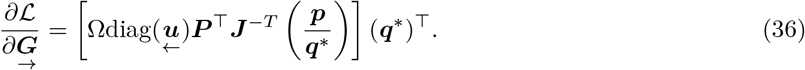

To optimise the parameters, we update 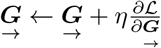, and project the result onto the set of lower-blocktriangular positive matrices. The corresponding expression for 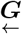 is obtained by symmetry of the forward and backward quantities.

## References

Bakermans, J. J., Warren, J., Whittington, J. C., and Behrens, T. E. (2023). Constructing future behaviour in the hippocampal formation through composition and replay. bioRxiv, pages 2023–04.

Basu, R., Gebauer, R., Herfurth, T., Kolb, S., Golipour, Z., Tchumatchenko, T., and Ito, H. T. (2021). The orbitofrontal cortex maps future navigational goals. Nature, 599(7885):449–452.

Ben-Yishai, R., Bar-Or, R. L., and Sompolinsky, H. (1995). Theory of orientation tuning in visual cortex. Proceedings of the National Academy of Sciences, 92(9):3844–3848.

Blanco-Pozo, M., Akam, T., and Walton, M. E. (2024). Dopamine-independent effect of rewards on choices through hidden-state inference. Nature Neuroscience, 27(2):286–297.

Botvinick, M. and Toussaint, M. (2012). Planning as inference. Trends in cognitive sciences, 16(10):485–488.

Botvinick, M. M. and Plaut, D. C. (2006). Short-term memory for serial order: a recurrent neural network model. Psychological review, 113(2):201.

Burak, Y. and Fiete, I. R. (2009). Accurate path integration in continuous attractor network models of grid cells. PLoS computational biology, 5(2):e1000291.

Burgess, P. W. and Wu, H. (2013). Rostral prefrontal cortex (Brodmann area 10). Principles of frontal lobe function, pages 524–544.

Callaway, F., van Opheusden, B., Gul, S., Das, P., Krueger, P. M., Griffiths, T. L., and Lieder, F. (2022). Rational use of cognitive resources in human planning. Nature Human Behaviour, 6(8):1112–1125.

Carlesimo, G. A., di Paola, M., Fadda, L., Caltagirone, C., and Costa, A. (2014). Prospective memory impairment and executive dysfunction in prefrontal lobe damaged patients: is there a causal relationship? Behavioural neurology, 2014(1):168496.

Chaudhuri, R., Gerçek, B., Pandey, B., Peyrache, A., and Fiete, I. (2019). The intrinsic attractor manifold and population dynamics of a canonical cognitive circuit across waking and sleep. Nature neuroscience, 22(9):1512–1520.

Chen, J., Zhang, C., Hu, P., Min, B., and Wang, L. (2024). Flexible control of sequence working memory in the macaque frontal cortex. Neuron.

Chiang, F.-K. and Wallis, J. D. (2018). Spatiotemporal encoding of search strategies by prefrontal neurons. Proceedings of the National Academy of Sciences, 115(19):5010–5015.

Clark, D. G., Abbott, L., and Sompolinsky, H. (2025). Symmetries and continuous attractors in disordered neural circuits. bioRxiv, pages 2025–01.

Dayan, P. (1993). Improving generalization for temporal difference learning: The successor representation. Neural computation, 5(4):613–624.

Donnarumma, F., Parr, T., Friston, K., Whittington, J., and Pezzulo, G. (2025). Inferential planning in the frontal cortex. bioRxiv, pages 2025–11.

Dorrell, W., Hsu, K., Hollingsworth, L., Lee, J. H., Wu, J., Finn, C., Latham, P. E., Behrens, T. E., and Whittington, J. C. (2024). Don’t cut corners: Exact conditions for modularity in biologically inspired representations. arXiv preprint arXiv:2410.06232.

Dorrell, W., Latham, P. E., Behrens, T. E., and Whittington, J. (2026). An efficient computing theory of prefrontal structured working memory representations. bioRxiv, pages 2026–02.

Duncan, J. (1990). Goal weighting and the choice of behaviour in a complex world. Ergonomics, 33(10–11):1265–1279.

Eckstein, M. K. and Collins, A. G. (2020). Computational evidence for hierarchically structured reinforcement learning in humans. Proceedings of the National Academy of Sciences, 117(47):29381–29389.

El-Gaby, M., Harris, A. L., Whittington, J. C., Dorrell, W., Bhomick, A., Walton, M. W., Akam, T., and Behrens, T. E. (2023). A cellular basis for mapping behavioural structure. bioRxiv, pages 2023–11.

Foster, D. J. (2017). Replay comes of age. Annu. Rev. Neurosci, 40(581–602):9.

Fuhs, M. C. and Touretzky, D. S. (2006). A spin glass model of path integration in rat medial entorhinal cortex. Journal of Neuroscience, 26(16):4266–4276.

George, D., Rikhye, R. V., Gothoskar, N., Guntupalli, J. S., Dedieu, A., and Lázaro-Gredilla, M. (2021). Clone-structured graph representations enable flexible learning and vicarious evaluation of cognitive maps. Nature communications, 12(1):2392.

Hafting, T., Fyhn, M., Molden, S., Moser, M.-B., and Moser, E. I. (2005). Microstructure of a spatial map in the entorhinal cortex. Nature, 436(7052):801–806.

Hopfield, J. J. (1982). Neural networks and physical systems with emergent collective computational abilities. Proceedings of the national academy of sciences, 79(8):2554–2558.

Iacaruso, M. F., Gasler, I. T., and Hofer, S. B. (2017). Synaptic organization of visual space in primary visual cortex. Nature, 547(7664):449–452.

Inagaki, H. K., Fontolan, L., Romani, S., and Svoboda, K. (2019). Discrete attractor dynamics underlies persistent activity in the frontal cortex. Nature, 566(7743):212–217.

Jensen, K. T. (2023). An introduction to reinforcement learning for neuroscience. arXiv preprint arXiv:2311.07315.

Jensen, K. T., Hennequin, G., and Mattar, M. G. (2024). A recurrent network model of planning explains hippocampal replay and human behavior. Nature neuroscience, 27(7):1340–1348.

Kim, S. S., Hermundstad, A. M., Romani, S., Abbott, L., and Jayaraman, V. (2019). Generation of stable heading representations in diverse visual scenes. Nature, 576(7785):126–131.

Kim, S. S., Rouault, H., Druckmann, S., and Jayaraman, V. (2017). Ring attractor dynamics in the drosophila central brain. Science, 356(6340):849–853.

Kingma, D. P. and Ba, J. (2015). Adam: A method for stochastic optimization. In Bengio, Y. and LeCun, Y., editors, 3rd International Conference on Learning Representations, ICLR 2015, San Diego, CA, USA, May 7-9, 2015, Conference Track Proceedings.

Kleinfeld, D. (1986). Sequential state generation by model neural networks. Proceedings of the National Academy of Sciences, 83(24):9469–9473.

Ko, H., Hofer, S. B., Pichler, B., Buchanan, K. A., Sjöström, P. J., and Mrsic-Flogel, T. D. (2011). Functional specificity of local synaptic connections in neocortical networks. Nature, 473(7345):87–91.

Kutschireiter, A., Basnak, M. A., Wilson, R. I., and Drugowitsch, J. (2023). Bayesian inference in ring attractor networks. Proceedings of the National Academy of Sciences, 120(9):e2210622120.

Lee, T. S. and Nguyen, M. (2001). Dynamics of subjective contour formation in the early visual cortex. Proceedings of the National Academy of Sciences, 98(4):1907–1911.

Levine, S. (2018). Reinforcement learning and control as probabilistic inference: Tutorial and review. arXiv preprint arXiv:1805.00909.

Mante, V., Sussillo, D., Shenoy, K. V., and Newsome, W. T. (2013). Context-dependent computation by recurrent dynamics in prefrontal cortex. nature, 503(7474):78–84.

Mattar, M. G. and Daw, N. D. (2018). Prioritized memory access explains planning and hippocampal replay. Nature neuroscience, 21(11):1609–1617.

Mattar, M. G. and Daw, N. D. (2026). Planning in the brain: It’s not what you think it is. Annual Review of Neuroscience, 49.

McNaughton, B. L., Battaglia, F. P., Jensen, O., Moser, E. I., and Moser, M.-B. (2006). Path integration and the neural basis of the’cognitive map’. Nature Reviews Neuroscience, 7(8):663–678.

Minka, T. P. (2013). Expectation propagation for approximate bayesian inference. arXiv preprint arXiv:1301.2294.

Mishchanchuk, K., Gregoriou, G., Qü, A., Kastler, A., Huys, Q. J., Wilbrecht, L., and MacAskill, A. F. (2024). Hidden state inference requires abstract contextual representations in the ventral hippocampus. Science, 386(6724):926–932.

Nyberg, N., Duvelle, É., Barry, C., and Spiers, H. J. (2022). Spatial goal coding in the hippocampal formation. Neuron, 110(3):394–422.

Ou, J., Qu, Y., Xu, Y., Xiao, Z., Behrens, T., and Liu, Y. (2025). Replay builds an efficient cognitive map offline to avoid computation online. bioRxiv, pages 2025–01.

Parmelee, C., Alvarez, J. L., Curto, C., and Morrison, K. (2022). Sequential attractors in combinatorial threshold-linear networks. SIAM journal on applied dynamical systems, 21(2):1597–1630.

Pedregosa, F., Varoquaux, G., Gramfort, A., Michel, V., Thirion, B., Grisel, O., Blondel, M., Prettenhofer, P., Weiss, R., Dubourg, V., et al. (2011). Scikit-learn: Machine learning in python. the Journal of machine Learning research, 12:2825–2830.

Ruff, D. A., Markman, S. K., Kim, J. Z., and Cohen, M. R. (2025). Linking neural population formatting to function. bioRxiv.

Schrit-twieser, J., Antonoglou, I., Hubert, T., Simonyan, K., Sifre, L., Schmitt, S., Guez, A., Lockhart, E., Hassabis, D., Graepel, T., et al. (2020). Mastering Atari, Go, chess and shogi by planning with a learned model. Nature, 588(7839):604–609.

Schultz, W., Dayan, P., and Montague, P. R. (1997). A neural substrate of prediction and reward. Science, 275(5306):1593–1599.

Shallice, T. and Burgess, P. W. (1991). Deficits in strategy application following frontal lobe damage in man. Brain, 114(2):727–741.

Shin, H., Ogando, M. B., Abdeladim, L., Durand, S., Belski, H., Cabasco, H., Loefler, H., Bawany, A., Hardcastle, B., Wilkes, J., et al. (2023). Recurrent pattern completion drives the neocortical representation of sensory inference. bioRxiv.

Skaggs, W., Knierim, J., Kudrimoti, H., and McNaughton, B. (1994). A model of the neural basis of the rat’s sense of direction. Advances in neural information processing systems, 7.

Solway, A. and Botvinick, M. M. (2012). Goaldirected decision making as probabilistic inference: a computational framework and potential neural correlates. Psychological review, 119(1):120.

Sompolinsky, H. and Kanter, I. (1986). Temporal association in asymmetric neural networks. Physical review letters, 57(22):2861.

Stachenfeld, K. L., Botvinick, M. M., and Gershman, S. J. (2017). The hippocampus as a predictive map. Nature neuroscience, 20(11):1643–1653.

Stroud, J. P., Watanabe, K., Suzuki, T., Stokes, M. G., and Lengyel, M. (2023). Optimal information loading into working memory explains dynamic coding in the prefrontal cortex. Proceedings of the National Academy of Sciences, 120(48):e2307991120.

Sutton, R. S. (1988). Learning to predict by the methods of temporal differences. Machine learning, 3:9–44.

Tian, Z., Chen, J., Zhang, C., Min, B., Xu, B., and Wang, L. (2024). Mental programming of spatial sequences in working memory in the macaque frontal cortex. Science, 385(6716):eadp6091.

Turner-Evans, D., Wegener, S., Rouault, H., Franconville, R., Wolff, T., Seelig, J. D., Druckmann, S., and Jayaraman, V. (2017). Angular velocity integration in a fly heading circuit. Elife, 6:e23496.

Turner-Evans, D. B., Jensen, K. T., Ali, S., Paterson, T., Sheridan, A., Ray, R. P., Wolff, T., Lauritzen, J. S., Rubin, G. M., Bock, D. D., et al. (2020). The neuroanatomical ultrastructure and function of a biological ring attractor. Neuron, 108(1):145–163.

van Opheusden, B., Kuperwajs, I., Galbiati, G., Bnaya, Z., Li, Y., and Ma, W. J. (2023). Expertise increases planning depth in human gameplay. Nature, pages 1–6.

Vinograd, A., Nair, A., Kim, J. H., Linderman, S. W., and Anderson, D. J. (2024). Causal evidence of a line attractor encoding an affective state. Nature, 634(8035):910–918.

Vollan, A. Z., Schellenberger, M. F., Moser, M.-B., and Moser, E. I. (2026). Attention-like regulation of theta sweeps in the brain’s spatial navigation circuit. bioRxiv, pages 2026–01.

Volle, E., Gonen-Yaacovi, G., de Lacy Costello, A., Gilbert, S. J., and Burgess, P. W. (2011). The role of rostral prefrontal cortex in prospective memory: a voxel-based lesion study. Neuropsychologia, 49(8):2185–2198.

Walton, M. E., Behrens, T. E., Buckley, M. J., Rudebeck, P. H., and Rushworth, M. F. (2010). Separable learning systems in the macaque brain and the role of orbitofrontal cortex in contingent learning. Neuron, 65(6):927–939.

Wang, J. X., Kurth-Nelson, Z., Kumaran, D., Tirumala, D., Soyer, H., Leibo, J. Z., Hassabis, D., and Botvinick, M. (2018). Prefrontal cortex as a meta-reinforcement learning system. Nature neuroscience, 21(6):860–868.

Wang, M., Fusi, S., and Stachenfeld, K. (2025). Brain-like slot representation for sequence working memory in recurrent neural networks. In Second Workshop on Representational Alignment at ICLR 2025.

Whittington, J. C., Dorrell, W., Behrens, T. E., Ganguli, S., and El-Gaby, M. (2023). On prefrontal working memory and hippocampal episodic memory: Unifying memories stored in weights and activation slots. bioRxiv, pages 2023–11.

Whittington, J. C., Muller, T. H., Mark, S., Chen, G., Barry, C., Burgess, N., and Behrens, T. E. (2020). The Tolman-Eichenbaum machine: Unifying space and relational memory through generalization in the hippocampal formation. Cell, 183(5):1249–1263.

Widloski, J. and Foster, D. J. (2022). Flexible rerouting of hippocampal replay sequences around changing barriers in the absence of global place field remapping. Neuron, 110(9):1547–1558.

Widloski, J., Theurel, D., and Foster, D. J. (2025). Spontaneous alternation of place-cell sequences in the open field through spike frequency adaptation. Cell Reports, 44(4).

Xie, Y., Hu, P., Li, J., Chen, J., Song, W., Wang, X.-J., Yang, T., Dehaene, S., Tang, S., Min, B., et al. (2022). Geometry of sequence working memory in macaque prefrontal cortex. Science, 375(6581):632–639.

Zhang, K. (1996). Representation of spatial orientation by the intrinsic dynamics of the headdirection cell ensemble: a theory. Journal of Neuroscience, 16(6):2112–2126.

Zheng, J., Guimaraes, R., Hu, J. Y., Perona, P., and Meister, M. (2024). Mice in the manhattan maze: Rapid learning, flexible routing and generalization, with and without cortex. Cognitive Computational Neuroscience.

Zintgraf, L., Shiarlis, K., Igl, M., Schulze, S., Gal, Y., Hofmann, K., and Whiteson, S. (2019). VariBAD: A very good method for Bayes-adaptive deep RL via meta-learning. arXiv preprint arXiv:1910.08348.

